# Cryptochrome-Timeless structure reveals circadian clock timing mechanisms

**DOI:** 10.1101/2022.07.27.501640

**Authors:** Changfan Lin, Shi Feng, Cristina C. DeOliveira, Brian R. Crane

## Abstract

Circadian rhythms influence many behaviors and diseases^1, 2^. They arise from oscillations in gene expression caused by repressor proteins that directly inhibit transcription of their own genes. The fly circadian clock offers a valuable model for studying these processes, wherein Timeless (TIM) plays a critical role in mediating nuclear entry of the transcriptional repressor Period (PER) and the photoreceptor Cryptochrome (CRY) entrains the clock by triggering TIM degradation in light^2, 3^. The cryo-EM structure of the CRY:TIM complex reveals how a light-sensing cryptochrome recognizes its target. CRY engages a continuous core of N-terminal TIM armadillo (ARM) repeats, resembling how photolyases recognize damaged DNA, and binds a C-terminal TIM helix reminiscent of the interactions between light-insensitive CRYs and their partners in mammals. The structure highlights how the CRY flavin cofactor undergoes conformational changes that couple to large-scale rearrangements at the molecular interface, and how a phosphorylated segment in TIM may impact clock period by regulating the binding of importin-α and the nuclear import of TIM:PER^4, 5^. Moreover, the structure reveals that the TIM N-terminus inserts into the restructured CRY pocket to replace the autoinhibitory C-terminal tail released by light, thereby providing a possible explanation for how the LS-TIM polymorphism adapts flies to different climates^6, 7^.

---

Circadian rhythms rely on cell-autonomous molecular clocks composed of transcriptional-translational feedback loops^1, 2^. In the paradigm clock of *Drosophila melanogaster* (**Figure 1a**), the transcriptional repressor proteins Period (PER) and Timeless (TIM) associate and accumulate in the cytosol before timed transport to the nucleus where they inhibit the transcription factors Clock (CLK) and Cycle (CYC)^2, 3^ and thereby downregulate the expression of *per*, *tim* and other clock-controlled genes^2, 3^ (**Figure 1a**). A cascade of TIM phosphorylation by the SHAGGY (SGG) and CK2 kinases gates nuclear entry of PER:TIM to properly time the circadian oscillator^4^. TIM co-transports PER by binding directly to the nuclear import factor importin-α1^5^.

**Figure 1.**
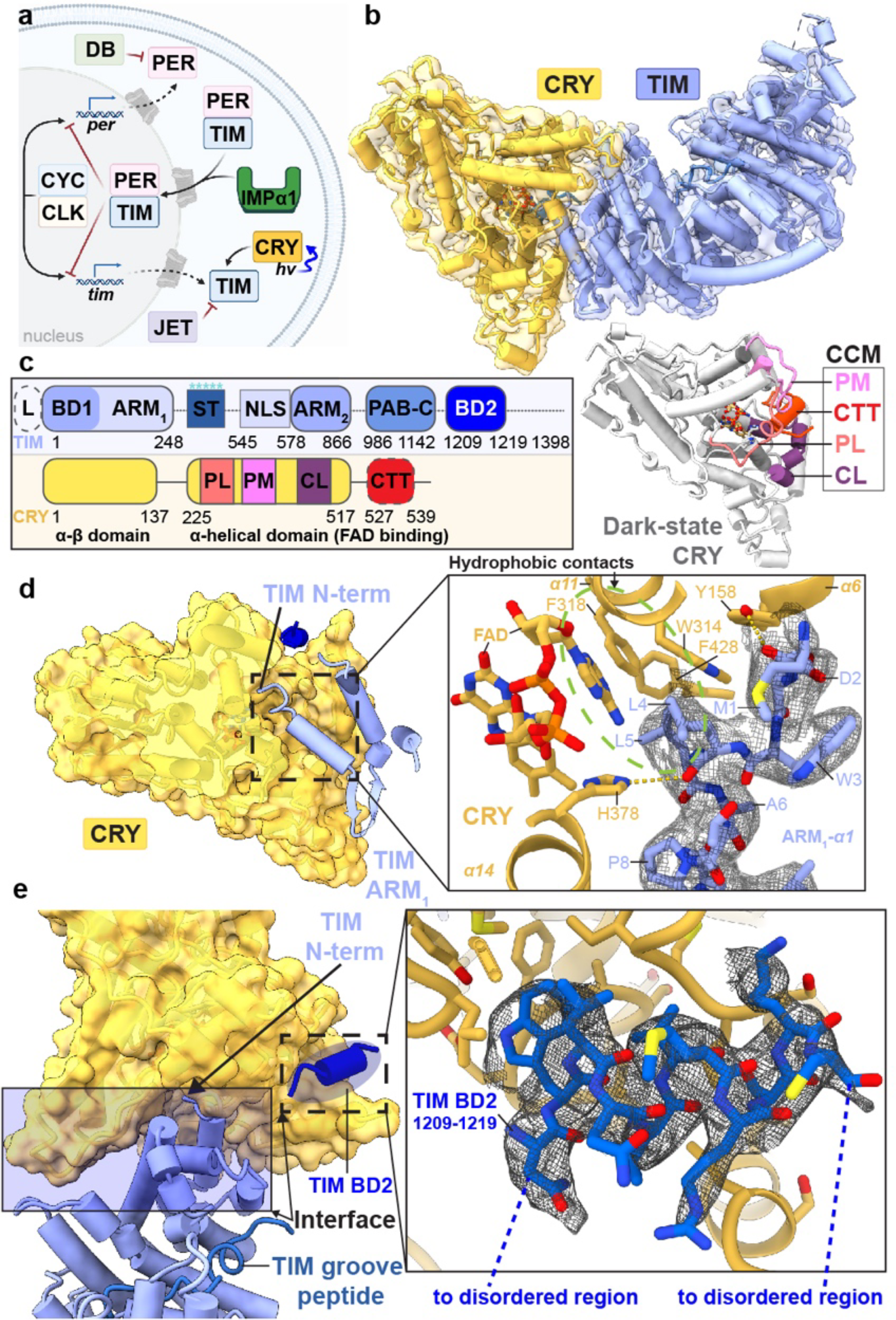
Cryo-EM structure of the CRY:TIM complex. (**a**) Drosophila Cryptochrome (CRY) entrains the circadian clock by initiating degradation of Timeless (TIM) in light. Jetlag (JET), Period (PER), Doubletime (DB), Cycle (CYC), Clock (CLK) and Importin-α1 (IMPα1) compose the core circadian oscillator. (**b**) CRY:TIM docked into the electron density map. Dark-state CRY shown at the bottom (grey, PDB ID: 4GU5). (**c**) Domain arrangements and key regions of CRY and TIM. Features of TIM: CRY-binding domains (CRY-BD1, 1-234, CRY-BD2, 1133-1398), serine/threonine-rich region (ST, 270-290) with key phosphorylation sites starred (S274, T278, T282, S286 and S290), ARM1 (1-248), ARM2 (578-866), nuclear localization sequence (NLS, 545-578), Parp-1 binding domain (PAB, 986-1142; TIM-C, 1051-1096), 23 residue L-TIM extension (L, (23aa)), disordered regions are dashed. Features of CRY: Phosphate binding loop (PL, 249-263), Protrusion motif (PM, 288-306), C-terminal lid (CL, 420-446), C-terminal tail (CTT, 519-542, absent in CRYΔ). (**d**) N-terminus of TIM inserts into the primary flavin binding pocket. 2.4 Å resolution local electron density is shown with notable hydrogen bonds (yellow dashes) and hydrophobic contacts (green circle). (**e**) Interface between CRY and TIM (blue shaded area), TIM groove peptide (in navy blue) and peripheral helix (fitted with 1209-1219 provisionally, blue). Inset shows the interactions between TIM 1209-1219 and CRY superimposed with electron density.

The fly clock is entrained by the flavoprotein Cryptochrome (CRY, **Figure 1a**), which binds to TIM in the light (**Figure 1b**) and directs it for proteasomal degradation^2, 3, 8^. CRYs share a flavin-binding Photolyase Homology Region (PHR) with the DNA repair enzymes photolyase (PL) but also contain a CRY C-terminal extension (CCE) that is specific for the function of a given CRY^9–11^. Type I CRYs from plants and animals sense light to reset the circadian clock; Type II CRYs from mammalians repress transcription in a light-independent manner by binding the PER proteins and directly inhibiting CLK^9, 10^. In Drosophila, photoreduction of the CRY FAD to the anionic semiquinone (ASQ) by a tryptophan tetrad (W342, W397, W420, W394) triggers release of a C-terminal tail (CTT) helix (**Figure 1b**)^12–15, 16–18^. Photoreduced CRY resets TIM levels by binding TIM and recruiting the E3 ubiquitin-ligase Jetlag (JET, **Figure 1a**)^8, 19, 20^. Recognition of TIM by CRY is hence the molecular event that sensitizes the clock to light. Although light-sensing CRYs have also been developed as powerful tools for optogenetic applications^21, 22^; there is little direct structural information on how light-activated CRYs recognize their targets.

## CRY:TIM structure determination

To overcome the challenge of isolating sufficient amounts of TIM (1398 residues) for structural characterization by cryo-electron microscopy (cryo-EM), we developed an insect (Drosophila S2) cell-based expression and nanobody-based affinity purification workflow for isolation of the 264 kDa CRY:TIM (**Supplementary Methods Overview**). To produce CRY:TIM, a CRY variant that lacks the 22-residue CCE^23^ and therefore binds TIM constitutively^24, 25^ was co-transfected and expressed in S2 cells with S-TIM. Co-expression with CRYΔ substantially increased TIM levels when ubiquitin-mediated degradation was also chemically blocked (see **Methods**). Nanobody-based affinity resin directed at a 13-residue ALFA tag allowed enrichment and elution of the complex. Complex homogeneity was further improved, and aggregation avoided by crosslinking with the lysine-specific reagent disuccinimidyl sulfoxide (DSSO) followed by SEC separation (**Extended Data Fig. 1**). Eluted samples were applied directly to glow-discharged Quantifoil® grids, vitrified, and imaged on a 200 keV Talos Arctica microscope equipped with a Gatan Quantum LS energy filter and K3 direct electron camera. Two out of three 3D classes were refined to yield global 3.48 Å (∼98 k particles) and 3.3 Å (∼160 k particles) maps of the complex **(Extended Data Fig. 1 and Supplementary Table 1**). We built an atomic model based on better resolved TIM density in the 3.3 Å map (**Supplementary Video 1**). Local resolution at the complex interface and within the body exceeds 2.5 Å.

## CRY recognizes the TIM N-terminus

The core of TIM revealed in the CRY:TIM complex (residues 1-248; 575-783; and 992-1128, **Figure 1b, Extended Data Fig. 2, and Supplementary Video 1)** forms a continuous right-handed supercoil of armadillo (ARM) 3-helix repeats^26^ in which two ARM domains discontinuous in sequence (ARM1 and ARM2, **Fig. 1c**) associate by swapping their terminal helices (**Extended Fig. 2**). TIM resembles mammalian TIM (mTIM, see below)^27^ but contains an N-terminal extension that binds to CRY **(Figure 1d**). The CRY PHR comprises an N-terminal α/μ domain and a C-terminal helical domain that harbors the FAD isoalloxazine ring in a central 4-helix bundle formed by α13-α16 (**Figure 1b,c, Extended Data Fig. 3, and Supplementary Video 1**)^28–30^. N-terminal to the bundle, helices α8-α11 compose the top of the flavin binding pocket and interact with the FAD phosphate groups and adenosine moiety (**Extended Data Fig. 3**). In this cofactor-capping region the phosphate binding loop (residues 249-263) following α8 and the protrusion motif (residues 288-306) following α10, distinguish CRYs from PLs (**Figure 1c**)^28^. The phosphate-binding loop, the protrusion motif, and the Ser-rich C-terminal lid (residues 420-446) together comprise the C-terminal coupled motif (CCM, **Figure 1c and Extended Data Fig. 3**) that changes proteolytic sensitivity when CRY becomes light activated^13^.

**Figure 2.**
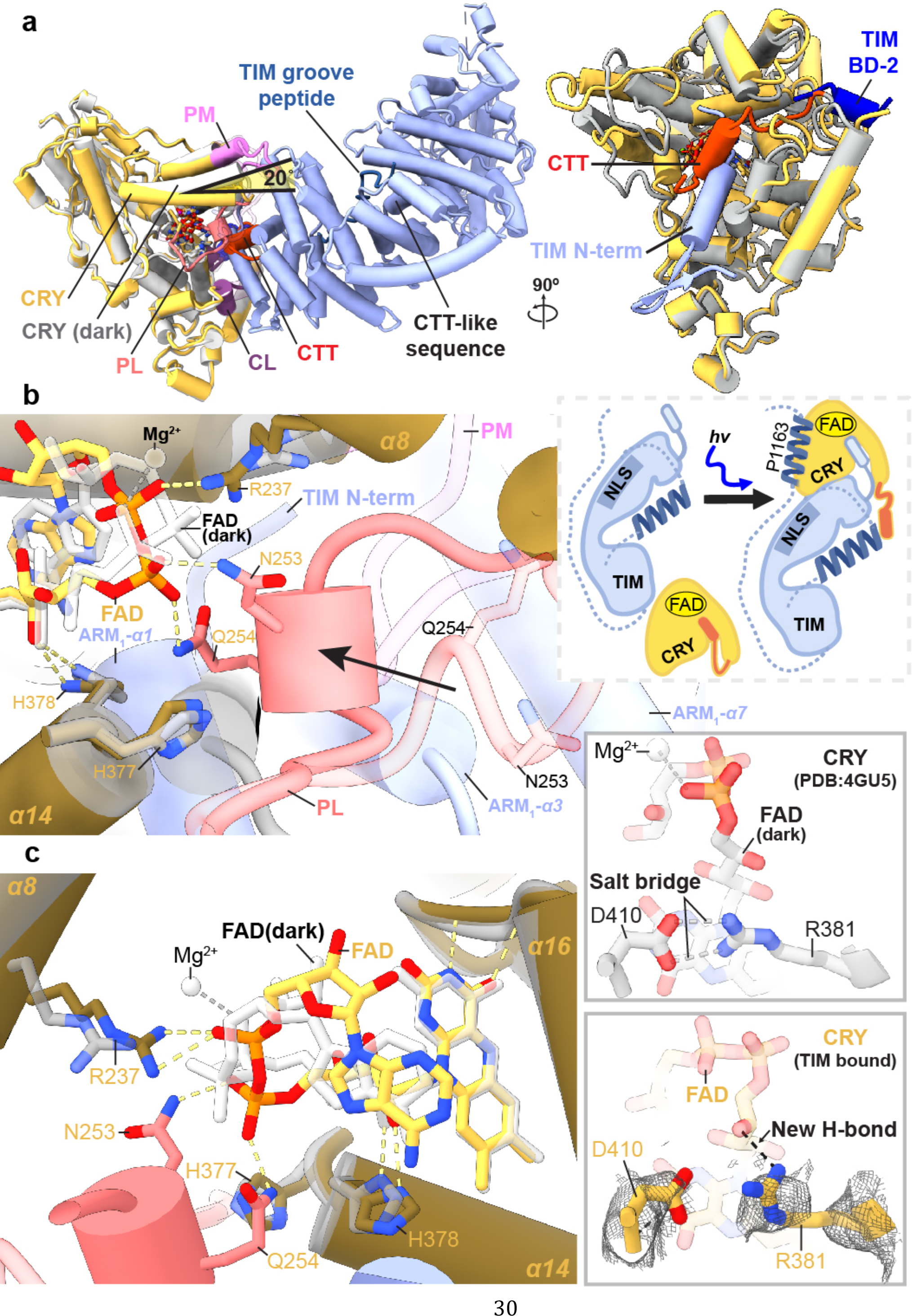
Conformational changes of FAD to the CRY C-terminal coupled motif upon binding TIM. (**a**) CRY:TIM (CRY, yellow; TIM, blue) in superposition with dark-state CRY. The N-terminus of TIM replaces the CRY CTT (red), as depicted in the schematic below. Structural elements designated as in Figure 1. A CTT-like sequence on TIM (617-626) that in isolation binds light-state CRY^13^ locates in the ARM core remote from the interface and is thus unlikely to be a binding determinant. (**b**) The phosphate binding loop (dark pink) collapses into the flavin pocket and N253 and E254 hydrogen bond (yellow dashes) with the FAD diphosphate conformation (red, orange) that differs substantially from that of the dark state (white bonds). (**c**) R237 replaces the Mg^2+^ counter ion and H377 anchors the new position of the loop. The D410- R381 salt bridge (top) appears to break in response to the new conformation of the ribose backbone (bottom). Reduced side-chain density of R381 suggests increased flexibility in the TIM complex.

**Figure 3.**
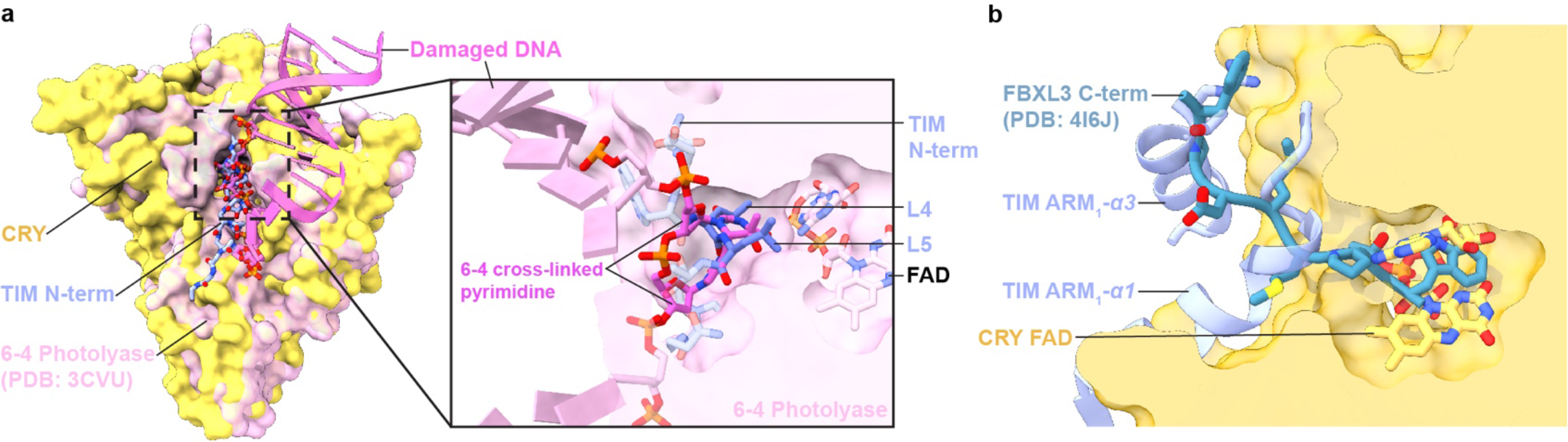
Conservation of recognition modes by CRY family members. (**a**) The interaction of CRY with TIM corresponds closely to how 6-4 photolyase recognizes the 6-4 pyrimidine lesion in damaged DNA (PDB ID:3CVU). TIM Leu4 and Leu5 superimpose on the crosslinked bases and the TIM ARM1- α1 polypeptide closely follows the path of 5’-to-3’ strand-separated DNA. (**b**) The TIM and FAD contacts of CRY mimic the interaction of apo-mCRY with the C- terminus of the FBXL3 E3 ligase (PDB ID: 4I6J).

TIM forms a discontinuous interface with CRY both at the N-terminus and C-terminus (**Figures 1b-e**). The principal CRY:TIM interface buries ∼1862 Å^2^ (49 residues) and 1807 Å^2^ (52 residues) of solvent accessible surface area on TIM and CRY, respectively. TIM primarily binds to CRY through the first three helices of the N-terminal ARM domain (**Figure 1d,e**). ARM1-α1 binds across CRY α17 and inserts directly into the flavin binding pocket replacing the CRY C-terminal tail (**Figures 1d, 2a, and Extended Data Fig. 4a)**. The contact is dominated by TIM ARM1-α1-3, ARM1-α7 and the ARM1-α9-10 loop which mesh against the CRY CCM (**Figure 1e and Extended Data Figs. 2 and 4**). At the center of the interface, Met1 and Asp2 extend out from α1 with Met1 packing against the Trp4 indole and embedding among several elements: Tyr250 and Leu251 of the CRY phosphate binding loop, Gly310 on α11, and Tyr158-Gln159 on the α5-α6 loop that extends down from the CRY α/ý domain (**Extended Data Figs. 3 and 5**). TIM Leu4 and Leu5 insert deep into the pocket adjacent to CRY Trp314 and the FAD adenine moiety. TIM ARM1-α2 contacts the CRY C-terminal lid and ARM1-α7 contacts the CRY protrusion motif, wherein the α10 helix lengthens to project the tip region toward the Met-rich ARM1-α9-α10 loop at the interface periphery (**Extended Data Figs. 4c,d**).

**Figure 4.**
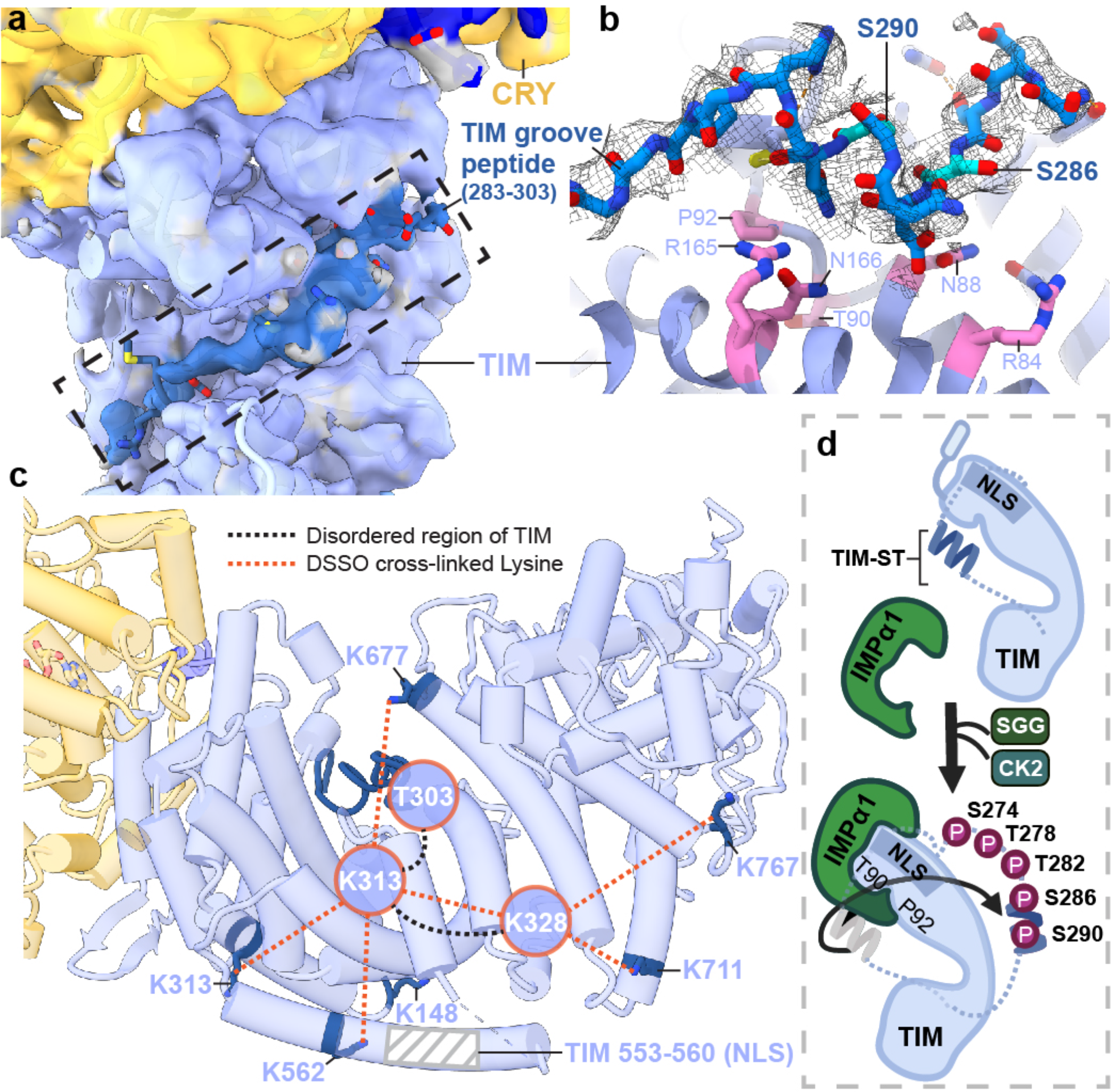
A structural basis for regulation of TIM nuclear entry. (**a**) Electron density (dark blue) for an extended polypeptide resides within a groove wherein members of the β-catenin and α-importin proteins bind their target sequences. (**b**) The TIM groove peptide density shown with the sequence assignment S283-T303. The peptide interacts with β-catenin-conserved Asn and Arg residues and buries residues important for importin-α1 binding (T90 and P92). (**c**) DSSO-mediated lysine crosslinking patterns of disordered residues K313 and K238 shown on the TIM structure. K313 and K328 produce a pattern of crosslinking that localizes them to a position consistent with placement of the S283-T303 groove peptide. The TIM NLS lies in a helix adjacent to the groove but does not occlude T90 and P92. (**d**) Schematic of proposed nuclear entry mechanism of TIM. Phosphorylation of the TIM-ST region by SGG and CK2 releases the ST region from TIM groove to expose TIM T90 and P92. Importin-α1 binds TIM adjacent to Thr90 and Pro92, while also binding the TIM NLS within a structurally analogous ARM groove of its own.

## CRY:TIM implications for photoadaptation

The recognition of TIM ARM1-α1 by CRY (**Figure 1d**) may explain an evolutionary adaptation in light-sensing by flies. A *Drosophila melanogaster tim* allele (*ls-tim*) causes differential expression of two isoforms; S-TIM, which begins at the N-terminal Met residue found in the structure, and L-TIM, which extends by an additional 23 N-terminal residues owing to introduction of an alternative translational start site (**Figure 1b**)^6,7^. The frequency of *ls-tim* correlates with geographical latitude and may confer a beneficial preference for more robust rhythms and a diapause state of dormancy in climates of more variable light and cooler temperatures. L-TIM reduces interaction with CRY by yeast two-hybrid, has greater stability in fly heads, and diminishes clock responses to light ^20, 31^. The Met1 amino-terminus projects into a solvent channel at the interface of the two proteins that could therefore accommodate the L-TIM extension (**Figure 1d and Extended Data Fig. 6a**). However, the recognition of S-TIM by CRY through the N-terminal Met residue will be altered by the L-TIM extension (**Figure 1d)**, thereby potentially explaining the capacity of *ls-tim* to alter fly light sensitivity.

## A discontinuous TIM binding element

In addition to the principal interaction between TIM ARM1 and CRY, strong electron density for a TIM helical segment is evident among the C-terminal lid, the C-terminal end of the last CRY PHR helix (α20) and the α5-α6 loop that projects down from the α/ý domain (**Figure 1e and Extended Data Fig. 3**). This TIM helix buries 496 Å^2^ of surface area, interacts with the light-state configuration of the C-terminal lid and binds at the position occupied by the residues that link α20 to the CTT in the dark-state structure, thereby supporting results from interaction studies^25^ and providing further evidence that the CTT undocks from the flavin pocket upon light activation^13, 15, 17^.

There are no connections of this density to the ordered TIM ARM core and thus its sequence assignment is provisional; however, after computational modeling of many predicted helical sequence registers, several gave reasonable agreement with the density and stereochemical constraints of the site; however, of these segments only 1209-1219 was also predicted to occupy the site by Alphafold2.1^32^ (**Extended Data Fig. 7a**). To further validate binding by the C-terminal disordered region, we performed the SWFTI assay (**Extended Data Fig. 8**) with TIM truncations and the CRY variant H377L that displays constitutive binding to TIM but also retains the CTT^16^. CRY bound TIM 1-1132 and 571-1398 similarly, but with reduced affinity compared to full-length S-TIM. TIM 1133-1398, which contains the proposed peripheral helix, alone bound to CRY-H377L (**Extended Data Fig. 8**). A known mutation of human TIM that removes the analogous region (R1081X) causes familial advanced sleep phase by reducing interactions between mammalian TIM and mCRY2^33^. Thus, although the roles of mCRY and CRY are not the same in animals and insects^2^, they may produce similar interactions with the otherwise disordered C-termini of their respective TIM proteins. These interactions may include competition with PER because TIM 1209-1219 associates with CRY in a region that coincides with where mCRY binds the mammalian Period proteins (**Extended Data Fig. 9a**)^34–36^.

## TIM proteins diverge in function

TIM and mTIM have similar structures in their respective ordered regions (**Extended Data Fig. 9b**), but their functions diverge. In fact, mTIM is more closely related to the Drosophila Timeout protein (TIM-2) than to TIM itself^27, 33^. Although mTIM and TIM-2 play some role in circadian rhythms neither are central constituents of the core mammalian oscillator^2^. In contrast, mTIM and its homologs (TOF1 in yeast, SWI1 in and TIM-1 in *C. elegans*) stabilize replication forks during DNA replication, pair sister chromatids and regulate S-phase checkpoints^37^. Tof-1 participates in the large replisome complex in which it binds directly to dsDNA (**Extended Data Fig. 9d**)^38^. TIM and mTIM share a C-terminal domain (PAB) that includes a conserved stretch of the Timeless C region (residues 1051-1096^33^) and binds poly-(ADP-ribose) polymerase – 1 (PARP-1) in mTIM^39^. However, just as the mTIM N-terminus is unsuited to bind mCRY, the TIM PAB is unlikely to bind to PARP-1 because it differs in the interface elements displayed by mTIM (**Extended Data Fig. 9c**). Thus, details of the respective structures indicate that the roles of TIM in the stabilization and co-transport of PER flies are likely distinct from those of mTIM, Tof-1, and TIM-2 in genome maintenance.

## Flavin conformational changes

Pronounced changes in the CRY cofactor binding pocket (**Figure 2a**) involve repositioning of the adenosine and ribose moieties, a large switch in the conformation of the diphosphate groups and refolding of the C-terminal lid to interact with TIM ARM1-α1 and α2. Moreover, the phosphate binding loop collapses into the flavin pocket to allow Asn253, Gln254 and Gln311 to hydrogen bond with the FAD phosphates (**Extended Data Fig. 5a)**. Tyr250 and Leu251 interact with the FAD pocket and make space for the top of TIM ARM1-α3 to insert Phe58 behind the new conformation of the phosphate-binding loop (**Figure 2b and Extended Data Fig. 5a**). In addition, cofactor-capping helix α8 tilts 20° towards the FAD diphosphates as Arg237 replaces the FAD Mg^2+^ counter ion, for which no electron density is observed (**Figure 2c and Extended Data Fig. 5b**). The FAD adenine ring slides 1.5 Å relative to the unbound structure, an adjustment compensated by a shift of Phe318 and harboring helix α11 (**Extended Data Fig. 5b**). The 262- to-264 loop also changes structure to accommodate the new diphosphate conformation and Gln311 swivels away from TIM Leu4 (**Extended Data Fig. 5a,c**). The primary light-sensitive reaction of flavin photoreduction and CTT release precedes TIM binding; however, the change in flavin redox state may induce the observed conformational changes of the FAD and serve to restructure CRY for downstream target engagement.

The CRY:TIM complex was purified under light. Although the cofactor redox state could not be verified on the cryo-EM grids, CRY light sensitivity studies^15^ indicate that the CRY flavin should be photochemically reduced under these conditions. Regardless, TIM likely stabilizes features of the light-state structure. In support of light-activated CRY in the complex, an Asp410-to-Arg381 salt bridge critical for flavin binding and the site of the light-insensitive *cryb* mutation^40^, appears to break in response to the new conformation of the ribose backbone (**Figure 2c**), in keeping with predictions of light activation from MD simulations^41^. Two conserved histidine residues within the flavin pocket, His377 and His378, participate in the PL catalytic mechanism and are implicated in CTT release and TIM binding^16, 18, 42^. In the unbound state His378 ND1 hydrogen bonds with O2’ of the FAD ribose, but in the TIM complex, His378 tilts away from the ribose hydroxyl to hydrogen bond to the TIM Leu5 carbonyl with NE2 (**Figures 2b,c and Extended Data Fig. 5b**). Substitutions at His378 still allow CTT release, but on/off switching is less robust^16, 42^. Protonation of His378 NE2 is likely required for it to interact with the TIM backbone carbonyl and His378 protonation may be responsive to the flavin redox state. The H377L substitution substantially increases the binding affinity to TIM in both light and dark^16^. Surprisingly, His377 does not contact TIM directly but rather anchors the restructured phosphate binding loop by hydrogen bonding to the Asn253 carbonyl (**Figure 2c**). CRY flavin reoxidation is reversible, as is redocking of the CTT^13, 15, 17^, which would complete with TIM for the flavin pocket and hence promote CRY release. However, clock resetting does not require complex dissociation because both TIM and CRY are degraded by the ubiquitin-proteosome pathway upon formation of their complex^8, 20^.

## Relationships among CRYs and photolyases

The interaction of CRY with TIM corresponds closely to how 6-4 photolyase recognizes the 6-4 crosslinked pyrimidine lesion in damaged DNA (**Figure 3a**). Indeed, TIM α1 residues Leu4 and Leu5 superimpose with the crosslinked base and the TIM polypeptide closely follows the path of 5’-to-3’ strand-separated DNA (**Figure 3a**)^43^. Key residues in the pocket also share identity and similar positions, with the notable exception of PL Tyr306, which as a Phe in CRY contacts the Leu5 side-chain. TIM and FAD also combine in CRY to mimic interactions of the FBXL3 E3 ligase with apo-mCRYs for their ubiquitin mediated degradation (**Figure 3b**)^44^. E3 ligase recruitment to CRY could only be similar if the flavin is released, which does not occur upon CRY:TIM association. More likely, the E3 ligases recognize structural features unique to the complex. Despite similarities at the primary cofactor pocket, the phosphate binding loop, protrusion motif and C-terminal lid differ in both sequence and conformation among PLs, mCRYs and CRY. CRY also contains a secondary pocket between its two domains that in PLs binds antenna cofactors and in mCRYs mediates contacts with other clock components^9, 45^. No cofactors or other constituents are found within the secondary pocket of the CRY:TIM structure; however, it composes a deep cleft typical of a recognition motif (**Extended Data Fig. 6b**).

## Regulation of TIM nuclear entry

The TIM structure provides insight into the gating of TIM:PER nuclear entry by the interaction between TIM and importin-α1 (IMPα1). TIM is homologous to the β-catenins, their related family members (HMP-2, p120, plakoglobin, plakophilin, and SYS-1) as well as the importin-α/β karyopherins involved in nuclear transport^5, 26, 46^. These ARM proteins conserve a positively charged groove on the concave side of their super-helical crescents where they bind extended peptide chains, such as nuclear localization signals (**Figure 4a,b and Extended Data Fig. 10**)^47, 48^. Two mutations known to block nuclear entry by reducing interactions between TIM and importin- α1 (Thr90Ala and Pro92Leu; S-TIM numbering)^5^ localize within the TIM groove (**Figure 4b**) but do not interact with the TIM nuclear localization sequence, which rather extends in a helix along the convex side of the protein (**Figure 4c**). An extended polypeptide with α-helical character binds into the TIM groove adjacent to catenin-conserved Arg (84, 165) and Asn (88, 166) residues (**Extended Data Fig. 10**), and Thr90/Pro92 (**Figure 4b**). Crosslinking mass spectrometry revealed that TIM Lys313 and Lys328 in the disordered TIM domain (249-545) react with Lys residues in the groove and toward the bottom of the crescent (**Figure 4c and Extended Data Fig. 10**), whereas Lys residues in other disordered regions (400-544; 784-991; 1129-1398) show little crosslinking to TIM or CRY. After extensive modeling, the sequence Ser283-Thr303 was provisionally assigned as the most reasonable fit to the polypeptide density (**Extended Data Fig. 7b**). This assignment places Asp287 and Asn291 adjacent to the conserved Asn/Arg residues and occludes the Thr90 and Pro92 residues involved in importin-α1 binding (**Figure 4b**).

Phosphorylation of the TIM-ST stretch of SGG and CK2 phosphorylation sites (SGG: S274, T278; CK2: T282, S286, S290; **Figures 1b and 4b**) increases nuclear entry rates of TIM in pacemaker neurons (s-LVs)^4^. The Ser283-Thr303 polypeptide bound within the TIM groove contains Ser286 and Ser290, which belong to this region of regulatory sites^4^. Thus, the structure suggests that phosphorylation at these sites displaces TIM-ST from the groove to expose the Thr90-Pro92 recognition motif for importin-α1 and thereby initiate nuclear import (**Figure 4d**). Thus, key delays that tune the clock period through nuclear entry likely depend on the interactions of a TIM autoinhibitory segment that alter upon its timed phosphorylation by circadian kinases.

The CRY:TIM structure reveals how a light-sensing cryptochrome engages its target. It also provides insight into how evolutionary relationships among the component proteins comprise regulatory circuits. Swapping the auxiliary CTT of CRY for the variable TIM N-terminus not only sets the circadian oscillator but also tunes its light sensitivity. In this interaction, the ability of TIM to serve as a peptide mimetic for a PL DNA substrate underscores how conserved structural features of PLs are also utilized by CRYs. In gating its partnership with CRY, TIM shares an autoinhibitory mechanism that resembles that which regulates importin-α^48–50^. The striking changes in FAD geometry that couple to CRY conformational rearrangement and TIM binding were unexpected. These new insights raise questions as to whether other flavoprotein light-sensors share similar mechanisms to convey light or redox signals from their cofactors.

## Methods

### Cloning

All DNA constructs (given in the Supplementary Information) were built by a modified Gibson assembly method and Q5 polymerase-based mutagenesis with home-made reaction mixture, as reported previously^51^.

### Transient co-transfection of S2 cells

S2 cells (cat# CRL-1963, ATCC) were maintained at 27 °C in 90% Schneider’s Drosophila Medium (Cat# 21720024, ThermoFisher) suppled with 10% heat-inactivated Fetal Bovine Serum (FBS, Cat# F4135, sigma), as reported previously^51^. pAC5.1 vectors of CLIP-CRYΔ and TIM- SNAP-HA-ALFA were delivered into S2 cells using the TransIT-Insect Transfection Reagent (Cat# MIR6100, Mirus) following a modified version of the manufacturer’s instruction. Briefly, about twenty 100 mm culture dishes (Cat# 353003, Corning) with 1000x10^4^ cells in 10 ml fresh growth medium were prepared on day 1. About 20 hours later, 10 μg DNA in total (CRYΔ: TIM mass ratio 1:1) were transfected by the TransIT-Insect reagent in each 100 mm dish. 72 hours after transfection, cells were harvested by centrifugation with 500 × g for 5 minutes at room temperature, washed once with chilled PBS and then stored at -80 °C until lysis.

### Nanobody-based affinity purification of the CRY:TIM complex

Two cycles of freeze-thaw were used to lyse the S2 cells as follows: S2 cell pellets from 200 ml of culture were resuspended with 16 ml lysis buffer (50 mM Tris, pH 8, NaCl 150 mM, 20% glycerol, 1x protease inhibitor (Cat# A32965, ThermoFisher)), divided as 800 μL aliquots in each tube and flash frozen in liquid nitrogen. The 800 μl aliquots were then quickly thawed in a room temperature water bath. The lysate aliquots were resupended (without vortexing) and the freeze-thaw cycle repeated. 100 U/ml benzonase (sc-202391, Santa Cruz Biotechnology) was added to the sample and incubated at 4 °C for 45 min to remove DNA/RNA contamination. The cell lysate was spun at 20000 x g for 20 minutes at 4 °C and further filtered with a 0.45 micron PVDF syringe filter (Cat# SLHV033NB, Millipore Sigma). 320 μL of ALFA resin 50% slurry (cat# N1512, Nanotag Biotechnologies) was equilibrated with lysis buffer and then incubated with the cell lysate at 4 °C for 1 h on an end-to-end rotator (Cat# 88881001, ThermoFisher scientific). The slurry was transferred to a 15 ml falcon tube, centrifuged at 1000 x g for 1 min and wash the resin with 7 mL wash buffer (HEPES 50 mM, pH 7.9, NaCl 150 mM, 10% glycerol). This process was repeated 3 times. Then the resin was resuspended in 1 mL wash buffer and split into two and applied to separate spin columns (Cat# 6572, BioVision), with 80 μL resin applied to each column. The spin columns were washed 4 times with 500 μl wash buffer and spun down at 1000 x g. 80 μL resin in each spin column was resuspended in 500 μL elution buffer (wash buffer with 200 μM ALFA peptide, cat# N1520, Nanotag Biotechnologies) and incubated in the cold room for 20 min on an end-to-end rotator. The columns were spun at 1000 x g for 1 min to collect the elution. The elution was repeated 3 times and all of the elution samples were then combined (3 mL in total).

### DSSO crosslinking coupled with SEC

The elution was concentrated to about 70 to 80 μL (while maintaining the final protein concentration at less than 2 mg/mL to ensure efficient crosslinking) with a 30 kDa vivaspin concentrator (Cat# 28932235, Cytiva). 50 mM stock solution of DSSO (cat# A33545,507 ThermoFisher or cat# 9002863, Cayman Chemicals) was prepared in dimethyl sulfoxide (DMSO). DSSO was added to the elution sample to a final concentration of 1 mM and incubated at room temperature for 45 minutes, followed by quenching for 5 min at room temperature with addition of 1 M Tris buffer (pH 8) to produce a final concentration of 20 mM Tris. The crosslinked complex was concentrated by 50 kDa vivaspin (Cat# 28932236, Cytiva) and purified by a Superose SEC column 6 10/300 (Cat# 29091596, Cytiva) or 3.2/300 (Cat# 29091598, Cytiva). The protein complex was eluted in SEC buffer (20 mM HEPES, pH 7.9, 150 mM NaCl) and concentrated to ∼40 μL by a 50 kDa cutoff vivaspin concentrator.

### Cross-Linking Mass Spectrometry (XL-MS)

SEC-purified crosslinked product was mixed with 4x Laemmli Sample Buffer (Cat# 1610747, BioRad) and 2.5% 2-mercaptoethanol (BME, Cat# 1610710, BioRad) and then boiled for 6 min. The sample was then loaded on an SDS-PAGE gel and stained with QC Colloidal Coomassie Stain (Cat #1610803, BioRad). The band of the complex (∼ 20 ug protein sample) was cut out and prepared for MS-MS analysis of crosslinked peptides as previously reported^52^ and described below.

### In-gel digestion of SDS gel bands

The protein band (∼ 20 µg protein) was sliced into ∼1 mm cubes. The excised gel pieces were washed consecutively with 400 μL deionized water followed by 50 mM ammonium bicarbonate, 50% acetonitrile and finally 100% acetonitrile. The dehydrated gel pieces were reduced with 200 μL of 10 mM DTT in 100 mM ammonium bicarbonate for 45 min at 56 °C, followed by alkylation with 200 μL of 55 mM iodoacetamide in 100 mM ammonium bicarbonate at room temperature in the dark for 45 min. Wash steps were repeated as described above. The gel was then dried and rehydrated with 100 μL LysC at 10 ng/µL in 50 mM ammonium bicarbonate containing 10% acetonitrile (ACN) and incubated at 37 °C for 3 h. After the 3 h incubation, 30 µL Trypsin at 43 ug/uL in 50 mM ammonium bicarbonate was added and the sample was continued to incubate at 37 °C for 18 h. The digested peptides were extracted from the gel twice with 200 μL of 50% acetonitrile containing 5% formic acid and once with 200 μL of 75% acetonitrile containing 5% formic acid. Extractions from each sample were pooled together and then divided equally in half.

One half was then filtered with 0.22 µm spin filter (Costar Spin-X from Corning), dried to dryness in the speed vacuum and analyzed as LysC/Trypsin digests. The second half was dried to dryness in the speed vacuum then reconstituted with 30 µL GluC at 33 ng/µL in 50mM ammonium bicarbonate (pH 7.9), incubated at 37 °C for 18 h. After incubation, formic acid was added to stop enzyme reaction. The digests were filtered with 0.22 μm spin filter, dried to dryness in the speed vacuum and analyzed as LysC/Trypsin/GluC digests. Each sample was reconstituted in 0.5% formic acid prior to LC- MS/MS analysis.

### Crosslinks identification by nanoLC-MS/MS

The digested product was characterized by NanoLC-MS/MS analysis at the Cornell Biotechnology Resource Center. The analysis was carried out using an Orbitrap Fusion^TM^ Tribrid^TM^ (Thermo- Fisher Scientific, San Jose, CA) mass spectrometer equipped with a nanospray Flex Ion Source, and coupled with a Dionex UltiMate 3000 RSLCnano system (Thermo, Sunnyvale, CA). Each sample was loaded onto a nano Viper PepMap C18 trapping column (5 µm, 100 µm × 20 mm, 100 Å, Thermo Fisher Scientific) at 20 μL/min flow rate for rapid sample loading. After 3 minutes, the valve switched to allow peptides to be separated on an Acclaim PepMap C18 nano column (2 µm, 75 µm x 25 cm, Thermo Fisher Scientific) at 35 °C in a 90 min gradient of 5% to 40% buffer B (98% ACN with 0.1% formic acid) at 300 nL/min. The Orbitrap Fusion was operated in positive ion mode with nano spray voltage set at 1.7 kV and source temperature at 275 °C. External calibrations for Fourier transform, ion-trap and quadrupole mass analyzers were performed prior to the analysis. Samples were analyzed using the CID-MS^2^-MS^3^ workflow, in which MS scan range was set to 375–1500 m/z and the resolution was set to 60,000. Precursor ions with charge states 3-8 were selected for CID MS^2^ acquisitions in Orbitrap analyzer with a resolution of 30,000 and an AGC target of 5 ×10^4^. The precursor isolation width was 1.6 m/z and the maximum injection time was 100 ms. The CID MS^2^ normalized collision energy was set to 25%. Targeted mass-difference triggered HCD-MS^3^ spectra were acquired in the ion trap with AGC target of 1 ×10^4^. When a unique mass difference (Δ=31.9721 Da) was observed in CID-MS^2^ spectra for ions with a single charge and multiple charges, the HCD collision energy at 38% and 28% was used respectively, for the subsequent MS^3^ scans. All data were acquired under Xcalibur 4.3 operation software and Orbitrap Fusion Tune application v3.4 (Thermo-Fisher Scientific).

### MS Data analysis

All MS, MS^2^ and MS^3^ raw spectra from each sample were searched using Proteome Discoverer 2.4 (Thermo-Fisher Scientific, San Jose, CA) with XlinkX v2.0 algorithm for identification of crosslinked peptides. The search parameters were as follow: four missed cleavage for either double digestion or triple digestion with fixed carbamidomethyl modification of cysteine, variable modifications of methionine oxidation, asparagine and glutamine deamidation, protein N-terminal acetylation. The peptide mass tolerance was 10 ppm, and MS^2^ and MS^3^ fragment mass tolerance was 20 ppm and 0.6 Da, respectively. The *Drosophila melanogaster* database with added targeted protein sequences was used for PD 2.4 database search with 1% FDR for report of crosslink results. Identified crosslinked peptides were filtered for Max. XlinkX Score > 40 containing at least two identified MS^3^ spectra for each pair of crosslinked peptides. Search Results were exported by the software as a spreadsheet. The crosslinked peptides were analyzed by xiNET for generating the viewer of inter or inner crosslinks between two target proteins.

### SWFTI assay

S2 cells were co-transfected as described above with pAC 5.1 vectors of CLIP-CRY-377L and TIM-SNAP-HA-ALFA variants at a mass ratio of 3:1. The pulldown assay was carried out as follows, similar to that previously described^51^. 5 mL of S2 cell pellets were lysed with 400 μL lysis buffer containing non-ionic detergent (50 mM Tris buffer, pH 8, 150 mM NaCl, 1% IGEPAL CA- 630 (Cat# I8896, Sigma Aldrich), 10% glycerol and 1x protease inhibitor (Cat# A32965, ThermoFisher)) was added to cell pellets harvested from 5 mL cells. EDTA was omitted to avoid disturbing SNAP/CLIP dye labeling. All buffer pHs were adjusted at the working temperature. After cells were incubated with lysis buffer on ice for 30 mins (flicking the tube every 10 min to avoid vortexing), cell debris was removed by spinning at 15,000 × g for 15 minutes at 4°C.

To label the lysate sample with SNAP/CLIP dyes, 1 part of lysate was diluted with 2 parts of labeling buffer (50 mM Tris buffer, pH 8, 150 mM NaCl, 10% glycerol) to decrease the concentration of non-ionic detergent to less than 0.5%, as recommended by the manufacturer (New England Biolabs). Then 27 μL of diluted lysate was mixed with 1 μL of 0.1 mM SNAP dye (Cat# S9102S, SNAP-Cell 647-SiR), 1 ul 0.1 mM CLIP dye (Cat# S9217S, CLIP-Cell 505) and 1 ul 30 mM DTT and incubated in the dark at room temperature for 1 hour. 10 uL 4x Laemmli sample buffer (Cat# 1610747, BioRad) and 1.5 μL 2-mercaptoethanol (BME) was added to the 30 μL mix, boiled at 95 °C for 5 minutes. After labeling with the SNAP/CLIP substrate, the sample was protected from bright light to avoid photobleaching. 20 μL of the sample was loaded on a gradient stain-free gel (Cat# 4568095, BioRad) to perform SDS-PAGE. The result was imaged with Chemidoc (BioRad) and quantified by image lab software (BioRad).

To label CRY and TIM on the affinity resin, ∼350 μL cell lysate was mixed with 10 μL pre-washed magnetic HA resin slurry (Cat# 88836, ThermoFisher) and incubated on an end-to-end rotator overnight at 4 °C. The dark sample was covered by foil and kept in the dark. Next day, the resin was washed with room temperature 500 μL TBS-T (20 mM Tris buffer, pH 7.6, 150 mM NaCl, 0.05% Tween-20) four times. Then HA resin was resuspended in 27 μL labeling buffer, 1 μL 0.1 mM SNAP dye, 1 μL 0.1 mM CLIP dye and 1 μL 30 mM DTT. The tube was incubated in the dark on an end-to-end rotator at room temperature for one hour. Subsequently, 10 μL 4x Laemelli sample buffer and 1.5 μL BME were added to the 30 μL resin mix. The tube was boiled at 95 °C for 10 minutes to denature and elute CRY and TIM from HA resin. SDS-PAGE and imaging were performed the same as above. Effective KD values were calculated as described elsewhere^51^.

### Negative Stain Electron Microscopy

Initial CRY:TIM samples were screened by negative-stain. Formvar/carbon 200 mesh copper grids (Electron Microscopy Sciences) was plasma cleaned by the GloQube glow discharge system (Electron Microscopy Sciences). Ten μL of 0.05 mg/ml CRY:TIM solution was applied onto the grid, incubated for 30 seconds, and the excess solution was blotted with filter paper. 10 μl of 2% uranyl acetate was then applied to the grid twice for 30-second incubation followed by blotting. The grid was air-dried for 5 minutes and visualized by Philips Morgagni 268 Transmission Electron Microscope at 100 keV.

### Cryo-EM grid preparation and data collection of CRY:TIM complex

Four μL of 0.5 mg/ml CRY:TIM solution was applied to glow-discharged grids (300 mesh Quantifoil Au, R1.2/1.3) that were purchased from Electron Microscopy Sciences. After 30 seconds incubation, the excess solution was blotted for 4 sec with filter paper by a Vitrobot Mark IV (Thermo Fischer Scientific). Grids were subsequently vitrified in liquid ethane and stored in liquid nitrogen. Cryo-EM images of CRY:TIM complex were collected on a Talos Arctica (Thermo Fischer Scientific) operated at 200 keV at a nominal magnification of 79,000x with Gatan GIF Quantum LS Imaging energy filter, using a Gatan K3 direct electron camera in counting mode, corresponding to a pixel size of 0.516 Å. A total of 1636 images stacks were obtained with a defocus rage of -0.8 to -2.0 μm by EPU (Thermo Fischer Scientific). Each stack movie was recorded for a total dose of 53 e-/Å^2^.

### Cryo-EM data processing of the CRY:TIM complex

Dose fractionated image stacks were subjected to motioncor2^53^ wrapped in Relion 3.1.3 ^54^ followed by patch CTF estimation in CryoSparc v3.2.0^55^. The micrographs whose CTF estimation is greater than 4 Å were chose for later processing, which yielded 1473 micrographs. Blob picker in CryoSparc was used on the three hundred randomly selected micrographs, which was followed by 2D classification and generation of ∼23k particles for the first-round Topaz train^56^. Then the trained model was applied to all 1473 micrographs and ∼48k particles were extracted and selected by 2D classification for the second-round of Topaz train particle identification. After two rounds of Topaz train and extraction, in totality ∼2M particles were picked. Following 2D classification non-relevant particles were removed and only the best classes were kept (∼350k particles). The particle stack was subjected to Bayesian polishing in RELION 3.1.3 and imported back to CryoSparc for the further Ab Initio reconstruction and Heterogenous refinement with 3 classes. The second and third classes were selected, and multiple rounds of Homogenous and Non-uniform Refinement were performed to yield a global 3.48 Å (∼98k particles) and a global 3.3 Å (∼160k particles) map using “gold standard” FSC at 0.143 cut-off, respectively. The 3.3 Å map is subjected to the model building. C1 symmetry was used throughout data processing. The workflow is provided in Extended Figure 1.

### Model building and refinement of the CRY:TIM complex

The initial CRY model is adapted from the PDB entry 4GU5, and the initial TIM model was fetched from AlphaFold Protein Structure Database entry P49021^57^. Both models were manually docked into the map by UCSF ChimeraX^58^. The model was subjected to ISOLDE^59^ to fit the loops which undergo conformational changes into the map density. Intrinsically disordered loops of TIM which were not visualized in the map were not included in the model. The CRY:TIM model was subjected to multiple rounds of automatic refinement using Phenix real space refinement, and manually building using Coot^60–62^. Validation was performed by MolProbity^63^. The final refinement statistics is provided in Table S1.

The unassigned polypeptide in the TIM grove was built in both directions and the sequence register slid along the backbone for a 150 residue stretch in the 250-400 range and each register real-spaced refined against the electron density and Q-score^64^ calculated. Given the helical nature of the density at one end of the groove, directionality of the sequence was best fit with the polypeptide having its N-terminal end closest to CRY and the C-terminal end running toward the Lys313/328 crosslinking partners at the base of the groove. Side chain density in the helical region showed a variable size pattern that was inconsistent with the repetitive patterns of much of the disordered region beyond residue 350. C-terminal to the groove peptide Lys313 and Lys328 are unresolved, but should be closely positioned to their crosslinking partners. To assign sequence to the disconnected TIM helix binding the PHR C-terminus secondary structure predictions on the regions of TIM not resolved in the ordered core were performed by RAPTORX^65^. All predicted helical sequences were fit into the density and refined in real-space with a range of sequence registers. As an additional check AlphaFold2.1^57^ was used to predict interactions between CRY and the disordered region of TIM, with the highest scoring predictions also tested for agreement with the electron density map. The disconnected helix displays strong side-chain density, which sets its directionality and ruled out many sequence placements. Given the resolution limitations of the cryo-EM maps, we consider the sequence assignments of both of these helical regions tentative.

### Analysis of TIM structural homologs

Structural comparisons and sequence alignments of TIM structural homologs were carried out with the DALI server^66^.

## Supporting information

Supplementary Movie 1. Overview of the CRY:TIM cryo-EM structure

mass-spectroscopy data1

mass-spectroscopy data2

mass-spectroscopy data3

mass-spectroscopy data4

## Reporting summary

Further information on research design is available in the Nature Reporting Summary linked to this article.

## Data availability

The cryo-EM density maps and the atomic model for TIM:CRY have been deposited in the EM Data Bank (EMDB: EMD-27335) and Protein Data Bank (PDB: 8DD7), respectively. The mass spectrometry proteomics data have been deposited to the ProteomeXchange Consortium via the PRIDE partner repository with the dataset identifier PXD034054. All other data is contained within the supplementary information. Reagents are available from the authors upon request.

## Code availability

No custom computer code was used in the study.

### Acknowledgements

We thank Qin Fu and Sheng Zhang of the Proteomics and Metabolomics Facility of Cornell University for providing the mass spectrometry data; Katherine Spoth and Mariena Silvestray-Ramos for help with EM instrumentation; Yitong Li for suggesting use of the ALFA Tag and experimental assistance, Greg Merz, Thuy-Tien Nguyen, and Cody Aplin for general aid; and Richard Cerione and Michael Young for helpful feedback. Figures 1A, inlet in 2B, 4D, and S1 were created with BioRender.com. This work was supported by NIH grants R35GM122535 to BRC and 1S10 OD017992-01 for the Orbitrap Fusion mass spectrometer. This work made use of the Cornell Center for Materials Research Shared Facilities which are supported through the NSF (DMR-1719875).

## Author contributions

All authors contributed to each aspect of the study, but primarily, C.L., and C.C.D developed the SWFTI assay, C.L. performed CRY:TIM binding assays, C.L. assisted by C.C.D. developed methods to produce the CRY:TIM complex, C.L. prepared the complex for structural analysis. S.F. prepared samples for cryo-EM, collected and processed cryo-EM data, S.F. and B.R.C. built the CRY:TIM model, C.L., S.F., and B.R.C. analyzed the structure, C.L., S.F., C.C.D, and B.R.C. wrote the manuscript, B.R.C. conceived of and oversaw the project.

## Corresponding author

Correspondence to Brian R. Crane

## Extended Data Figure Legends

**Extended Data Figure 1.**
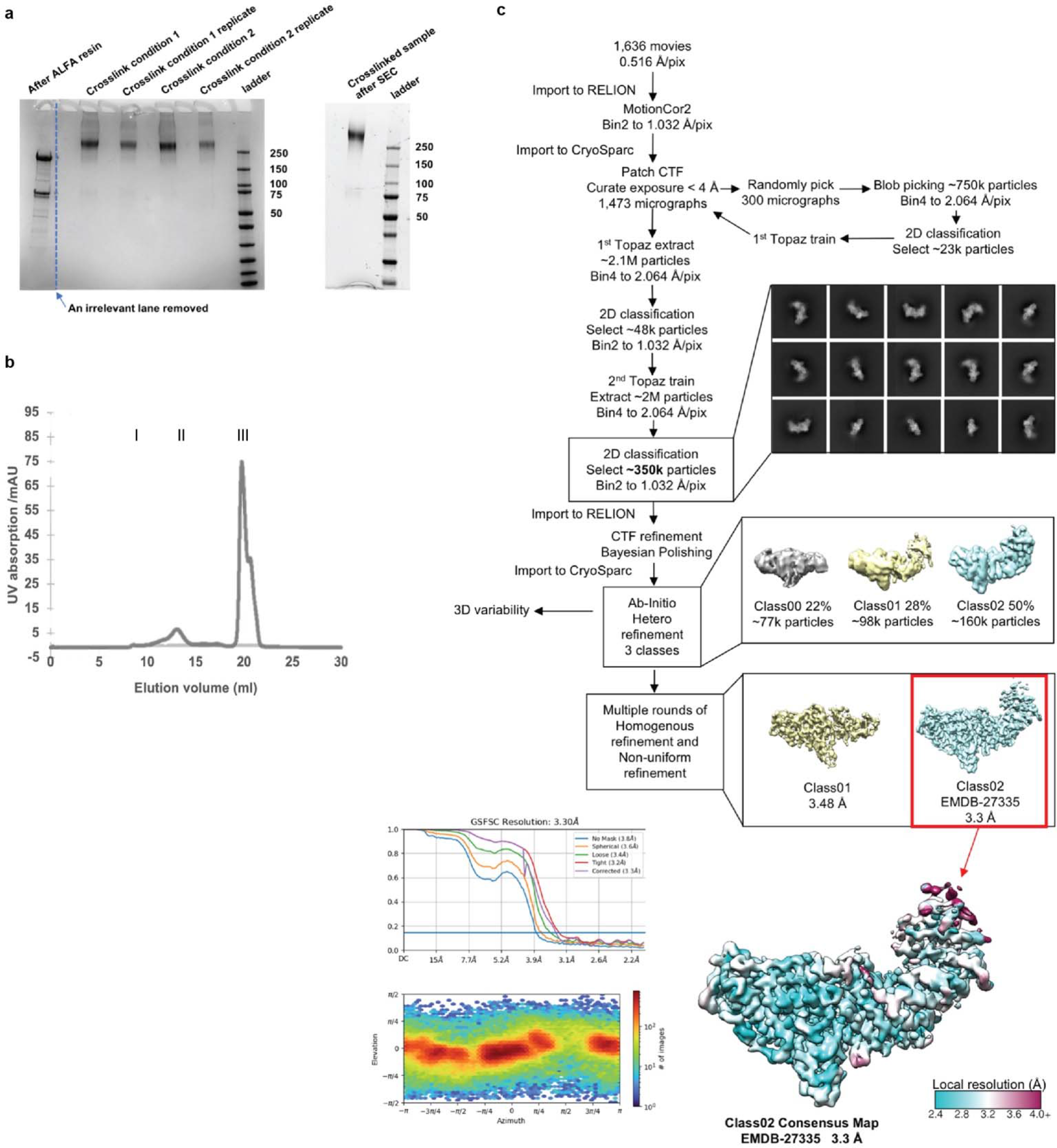
CRY:TIM purification and cryo-EM workflow. Purification of CRY:TIM complexes for cryo-EM. (a) SDS-PAGE gel showing complex after elution from the ALFA resin, followed by crosslinking, followed by SEC. (b) SEC trace of purified complex, peak I: aggregates, peak II: CRY-TIM complex, and peak III: excessive ALFA peptide. (c) Processing workflow for CRY:TIM collected on a Talos Arctica (Thermo Fischer Scientific) operated at 200 keV using a Gatan K3 direct electron camera. The red box indicates the final reconstruction in this study. Data processing was carried out in CryoSparc and RELION. Motion Correlation was performed by RELION (MotionCor2); with Contrast Transfer Function (CTF) applied in CryoSparc. Topaz was used for particle classification. Overall resolution determination in CryoSparc by Gold Standard Fourier Shell Correlation (GSFSC) is shown at the mid-bottom left. Corrected values give 3.30 Å overall resolution. Local resolution exceeds 2.5 Å, as shown on the bottom right. Particle orientational sampling is reasonably uniform as shown at the bottom left. Number of particles (images) holding a given azimuthal and elevation angle are indicated in the 2- dimensional histogram.

**Extended Data Figure 2.**
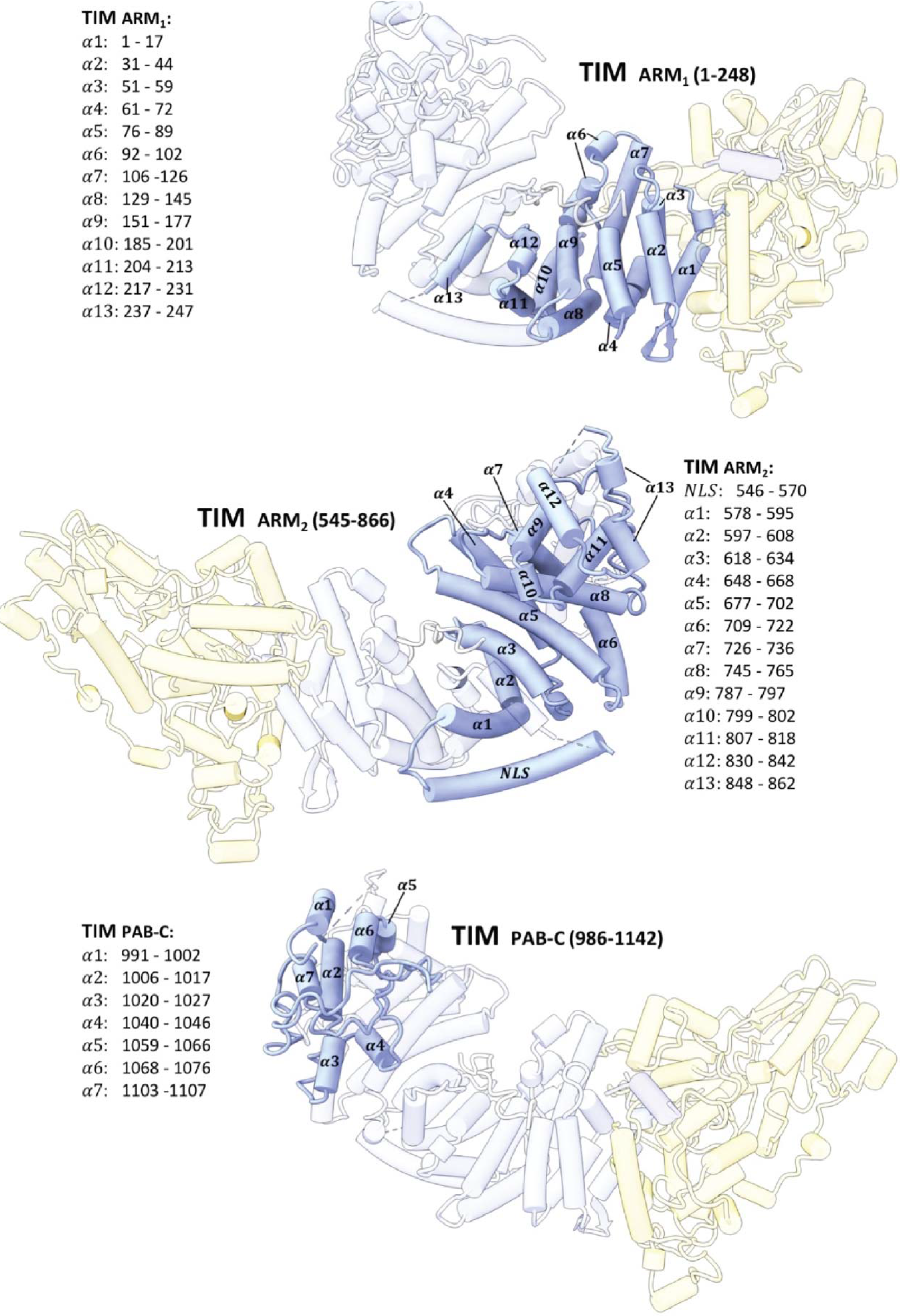
**TIM domain construction and secondary structure assignments**. Domain construction and secondary structure assignments of TIM.

**Extended Data Figure 3.**
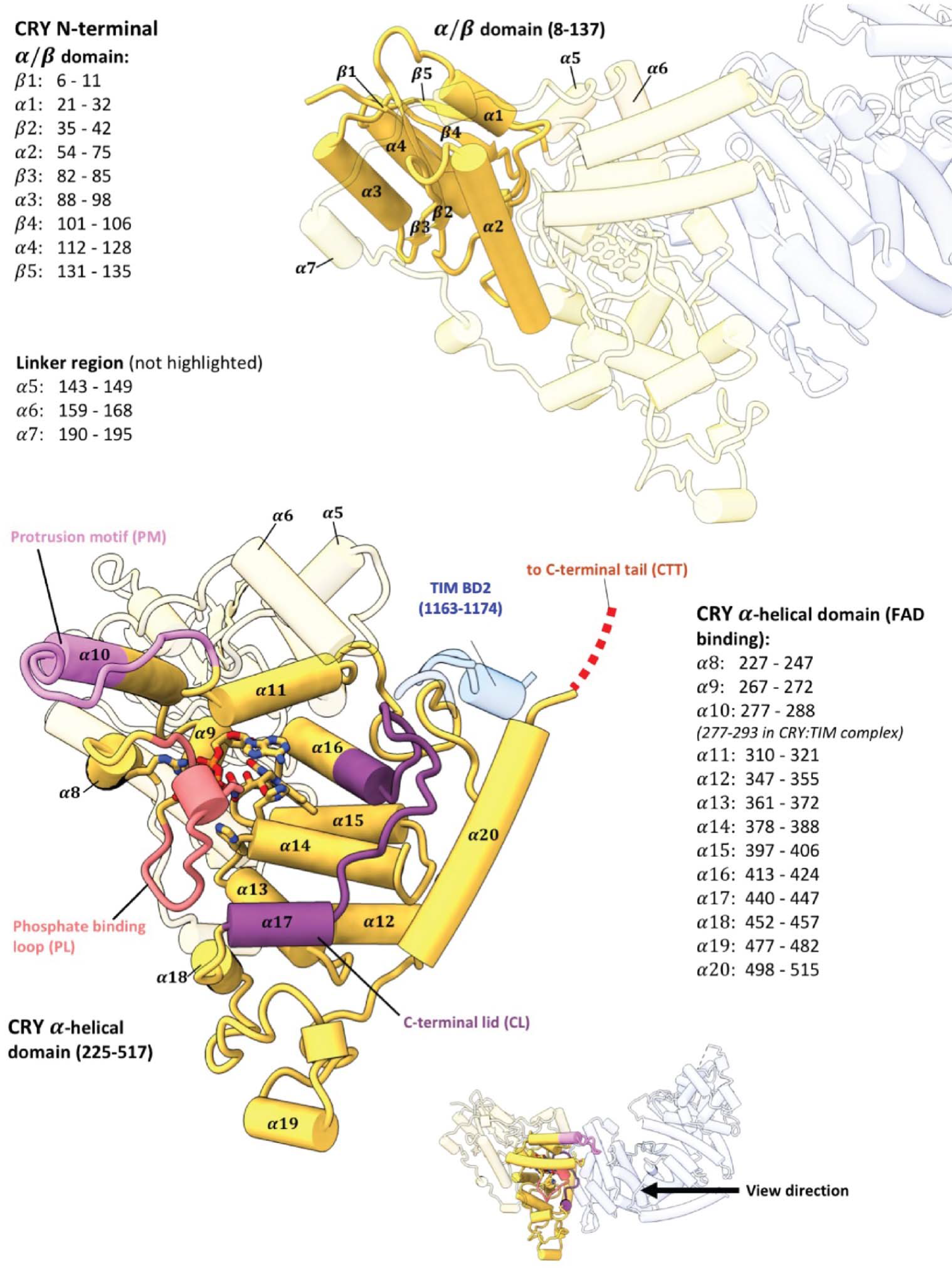
CRY domain construction and secondary structure assignments. Domain construction and secondary structure assignments of CRY.

**Extended Data Figure 4.**
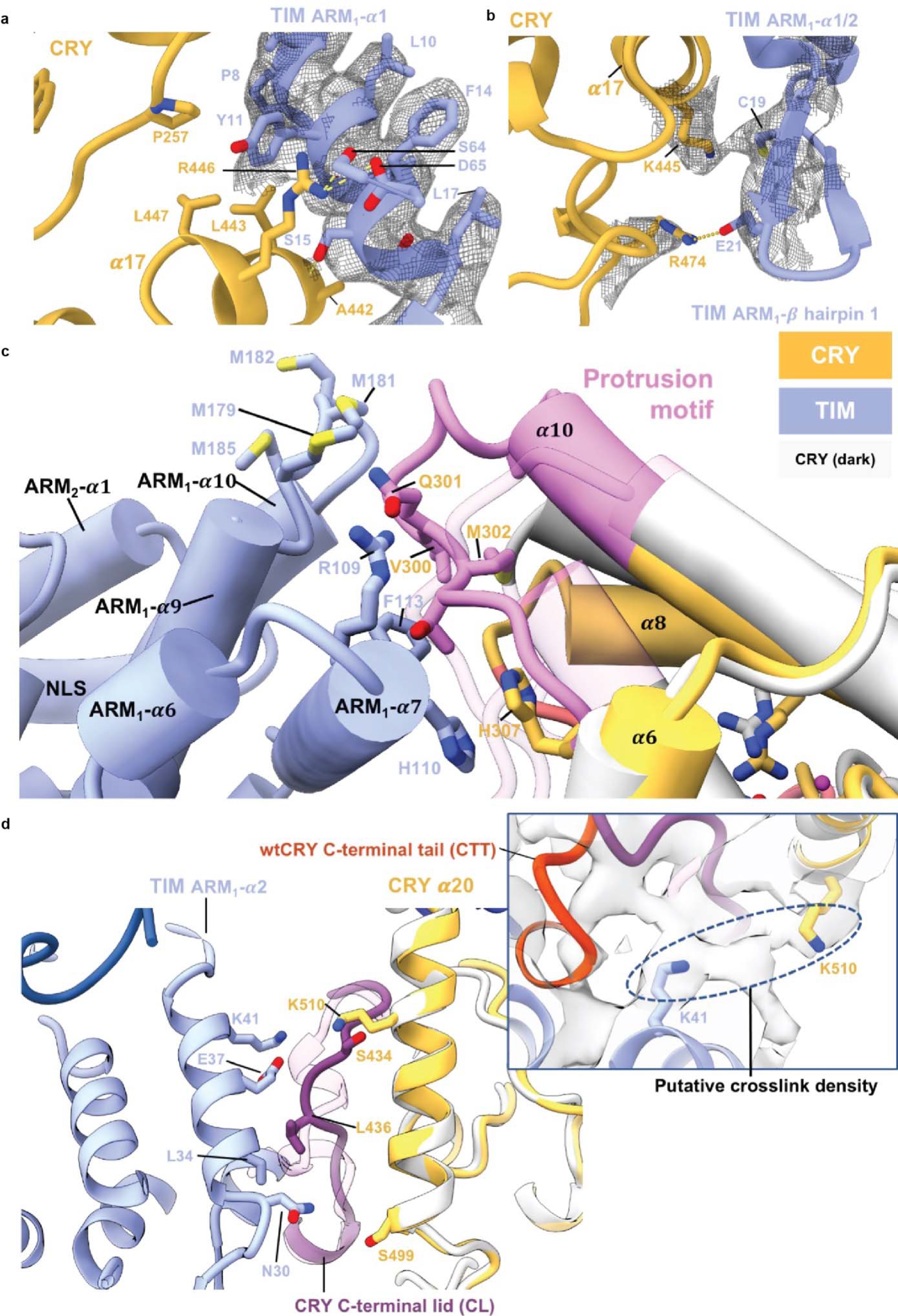
Key interactions at the CRY:TIM interface. (a) Interactions between the bottom half of TIM ARM1-α1 with CRY. ARM1- α1 binds across CRY α14 and α17. The guanidinium of CRY R446 (on α17) forms π-cation interaction with TIM Y11. TIM Y11 participates in hydrophobic contacts with CRY P257, L443 and L447. On TIM α1, TIM P8 stacks against CRY α12 and TIM S15 hydrogen bonds to the backbone carbonyl of CRY A442. (b) In the β-hairpin connecting ARM1- α1 to ARM1- α2 E21 salt-bridges to R474 and C19 interacts with K445. (c) Interactions of the CRY protrusion motif. TIM R109, F113, and H110 interact with the protrusion motif of CRY. TIM ARM2-α3 H110 stacks with CRY H307. Four Met residues (M179, M181 M182, M185) in the ARM3-α2-α3 loop provide side-chain interactions with the tip of the CRY protrusion motif. The CRY α10 helix lengthens to residue A295 compared to the dark state and the following 300-306 residues interact directly with ARM2-α3. (d) Interactions of the CRY C-terminal lid and α20 helix. The restructured C-terminal lid interacts with residues on TIM ARM1- α2. For examples, L34 make a hydrophobic contact with I436 and E37 hydrogen bonds to the backbone of S434. At the end of ARM1- α2, Asn30 contacts the N-terminal end of CRY α20. A DSSO crosslink observed between Lys510 on CRY a20 and Lys41 on ARM1-α3 is fully consistent with the observed interface.

**Extended Data Figure 5.**
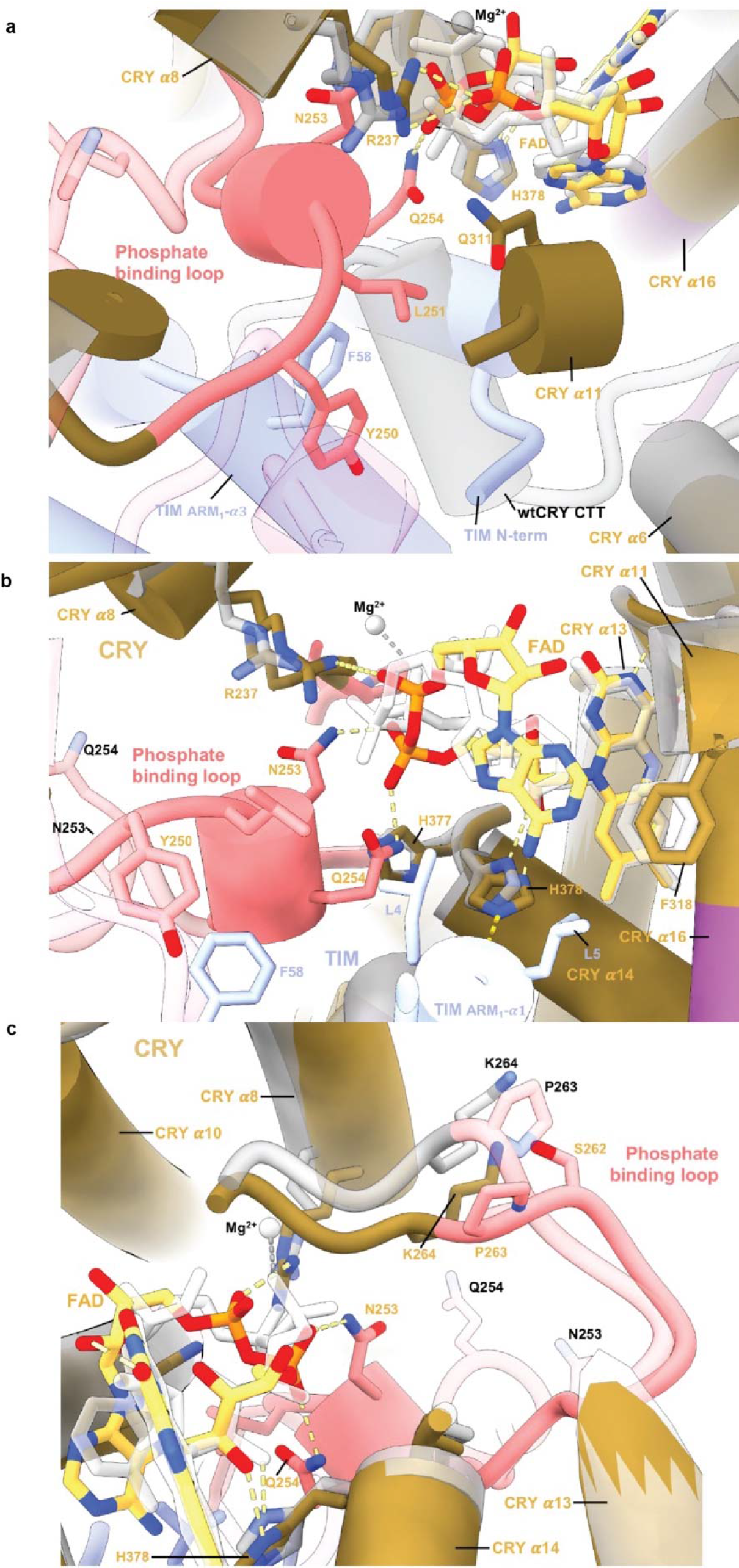
Changes in flavin binding pocket for CRY dark and bound states. Comparison of the flavin pocket in CRY:TIM compared to in dark-state CRY (PDB 4GU5). (a) In CRY:TIM, Y250 and L251 interact with the FAD pocket and make space for the top of TIM ARM1-α3 to insert F58 behind the new conformation of the phosphate-binding loop. (b) The FAD adenine ring of CRY:TIM slides 1.5 Å relative to the unbound structure. CRY F318 shifts away from its original position and harbors helix α11 to compensate for the FAD adenine ring adjustment, following the protrusion motif. (c) G311 swivels away from the inserted TIM L4. New diphosphate conformations cause rearrangement of the CRY 262-to-264 loop.

**Extended Data Figure 6.**
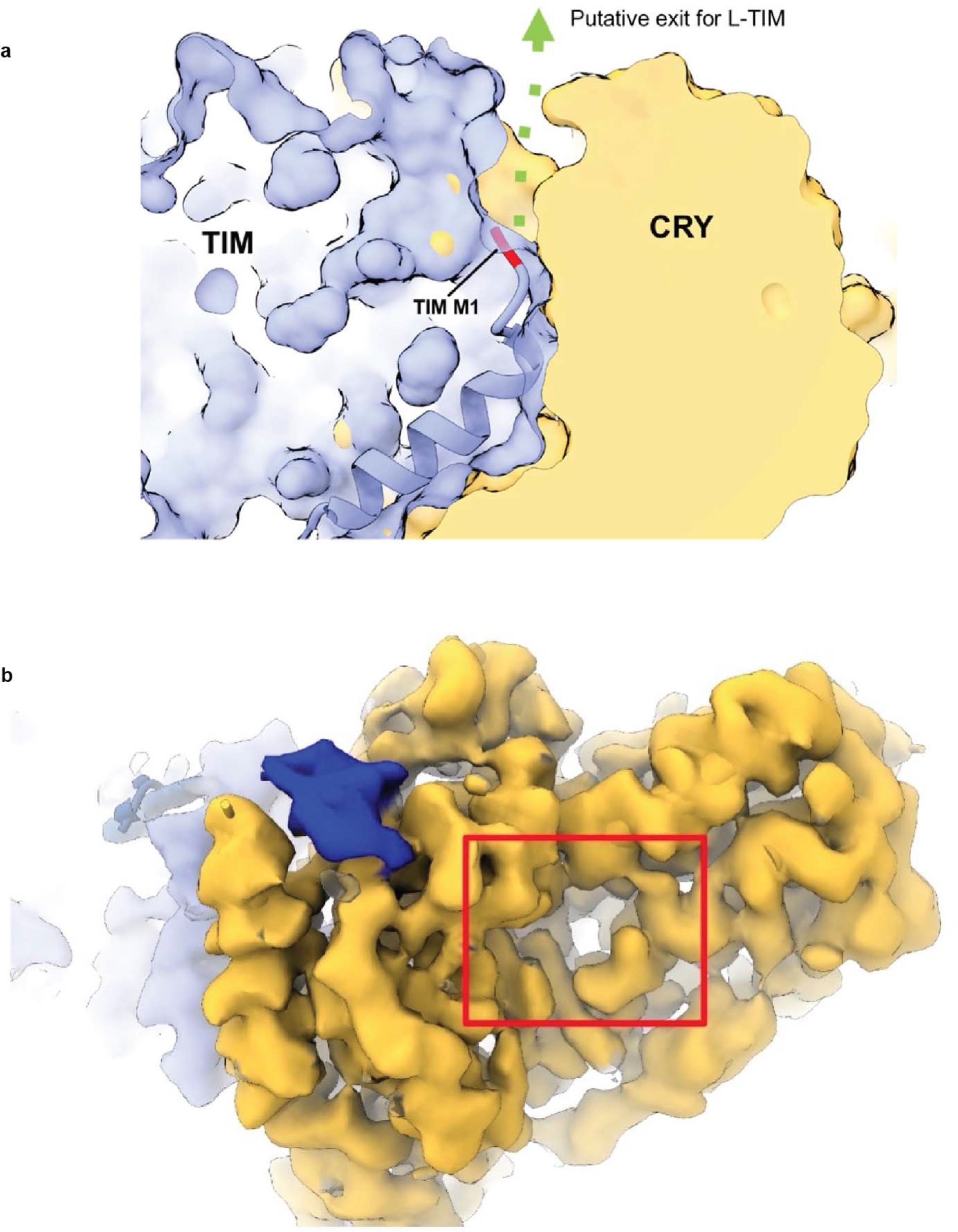
Exit channel at the CRY:TIM interface and CRY secondary pocket. (a) Putative exit channel for the additional 23 N-terminal residues of L-TIM. The first residue of TIM (M1) is colored as red. A solvent accessible channel at the interface between the two proteins is marked as the green arrow. (b) Electron density for the secondary binding pocket of CRY where antenna cofactors bind in photolyases and mCRY interact with other Clock proteins (**red** box) CRY (**yellow**), TIM (**periwinkle blue**), TIM 1163-1174 (**blue**), TIM groove (**navy blue**). Although the pocket is well formed, there is no obvious density for additional moieties.

**Extended Data Figure 7.**
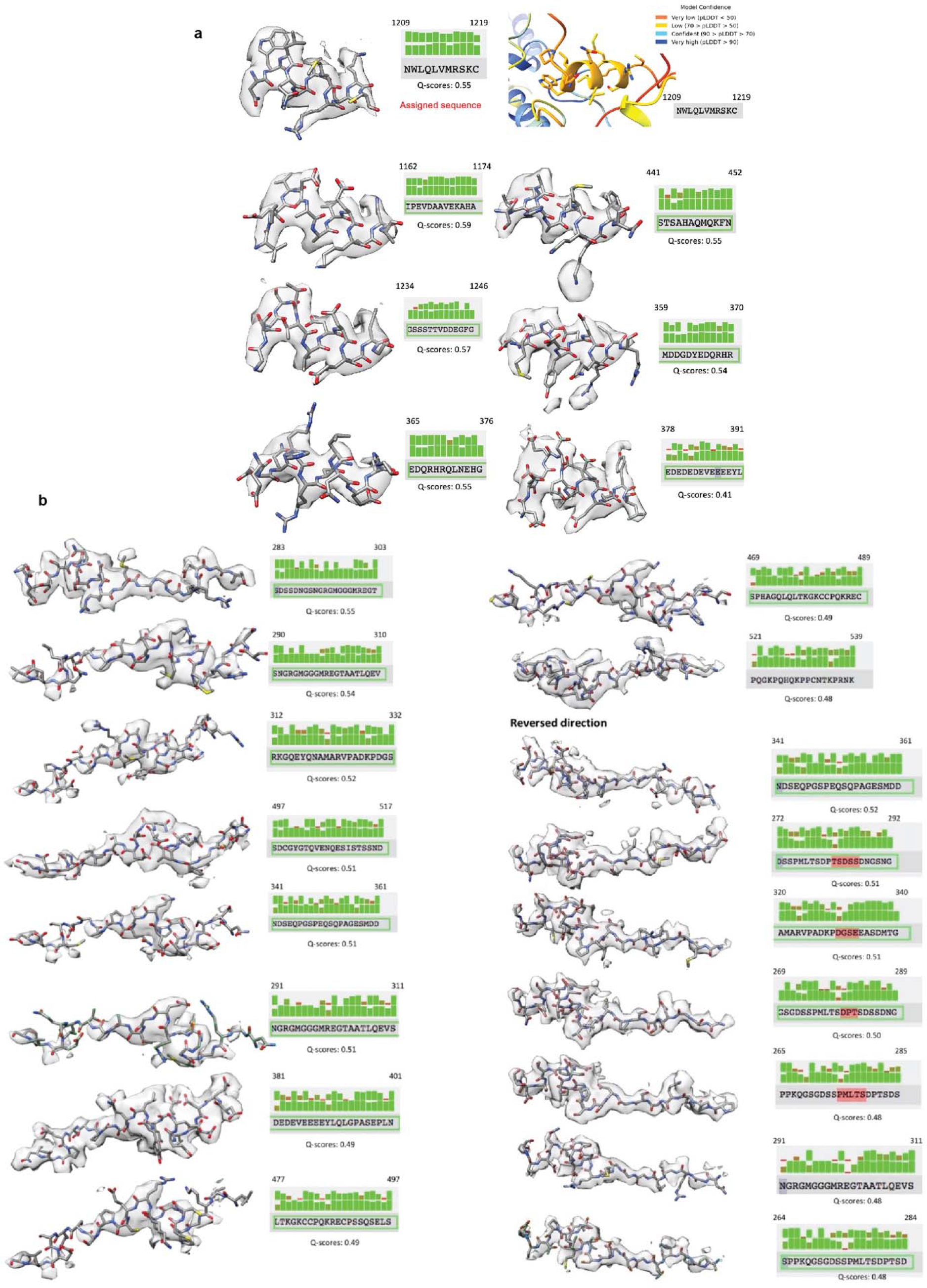
TIM peripheral helix and groove peptide fitting to the electron density map and associated Q-scores. (a) Real-space electron density fitting metrics for the top scoring sequence assignments for the TIM peripheral helix. Real space density agreements were calculated in Q-scores and are shown above for sidechains and below for mainchains. One of the top scoring segments, residues 1209-1219 (top left) was also predicted by AlphaFold2.1 to interact with CRY at the position of the observed electron density, the model confidence pLDDT scores for this segment are given at the top right. (b) Real-space electron density fitting metrics for the top scoring sequence assignments for the TIM groove-binding polypeptide. Real space density agreements were calculated in Q-scores and are shown above for sidechains and below for mainchains. The TIM groove binding polypetide was tested in both directions.

**Extended Data Figure 8.**
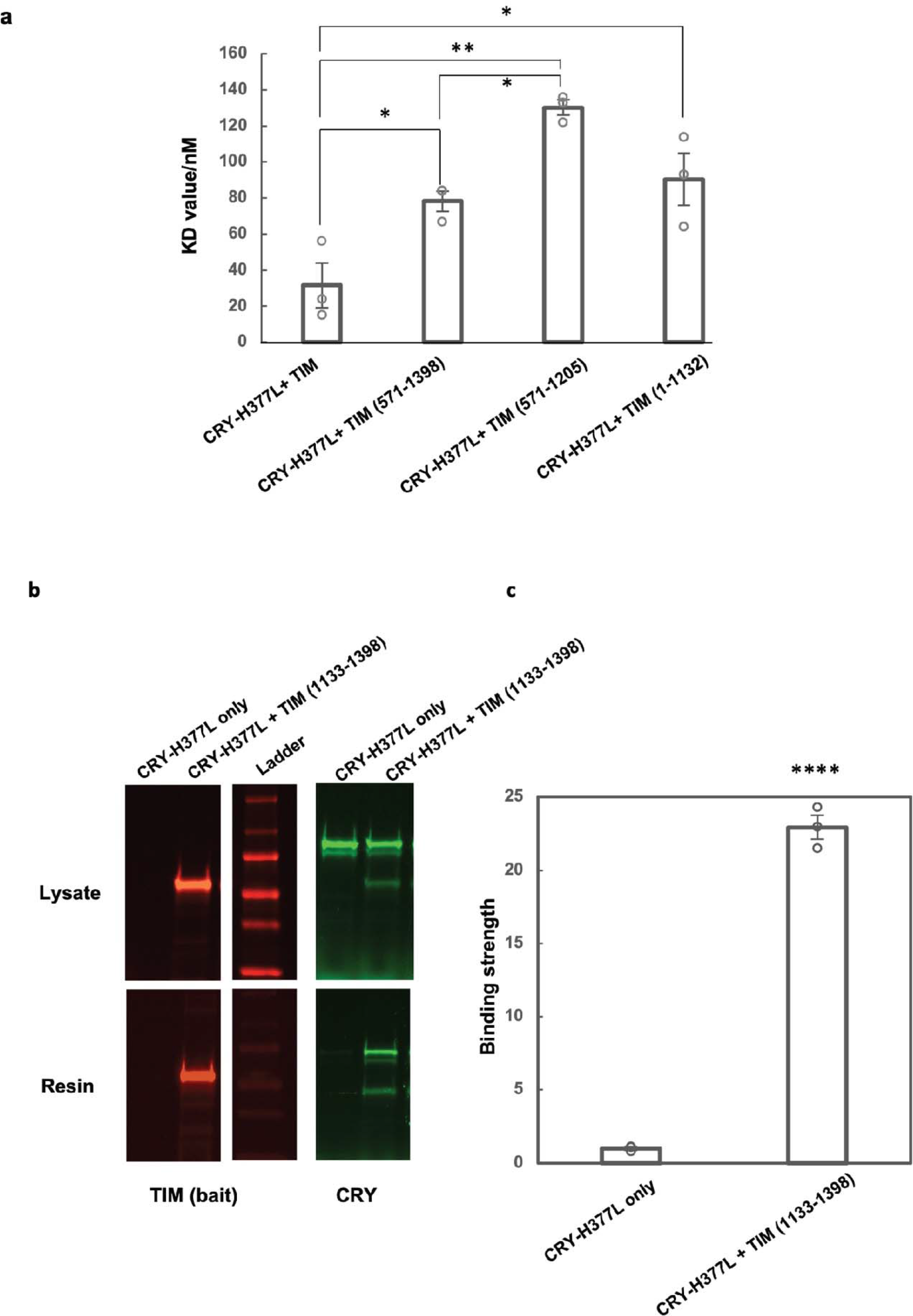
Relative binding affinities between fragments of TIM and CRY H377L by the SWFTI assay. (a) The CRY H377L variant acts as a constitutive light-activated state and enhances binding affinity for TIM by stabilizing the TIM-binding conformation of CRY^16^. H377L thereby increases TIM expression levels and facilitates detection of weaker binding variants. Mean ± standard error is shown for sample size = 3, from different biological replicates. One-way ANOVA with post-hoc Tukey HSD test was used to determine p values. (b) Pull down results of CRY-H377L with TIM-1133-1398. CRY-H377L interacts with the C-terminal regions of TIM. Gel lanes containing replicates or unrelated samples are not shown. (c) Quantification of the binding strength between CRY-H377L and TIM (1133-1398) compared to the negative control (CRY-H377L only with the same amount of resin). The binding strength is defined as the amount of CRY on the resin divided by the amount of CRY in the lysate sample. The negative control is normalized to 1. Mean ± standard error is shown for sample size = 3, from different biological replicates. Two-tailed unpaired t-test was used to determine p value. Binding affinities could not be derived from this experiment owing to the challenge of producing excessive CRY relative to TIM 1133-1398.

**Extended Data Figure 9.**
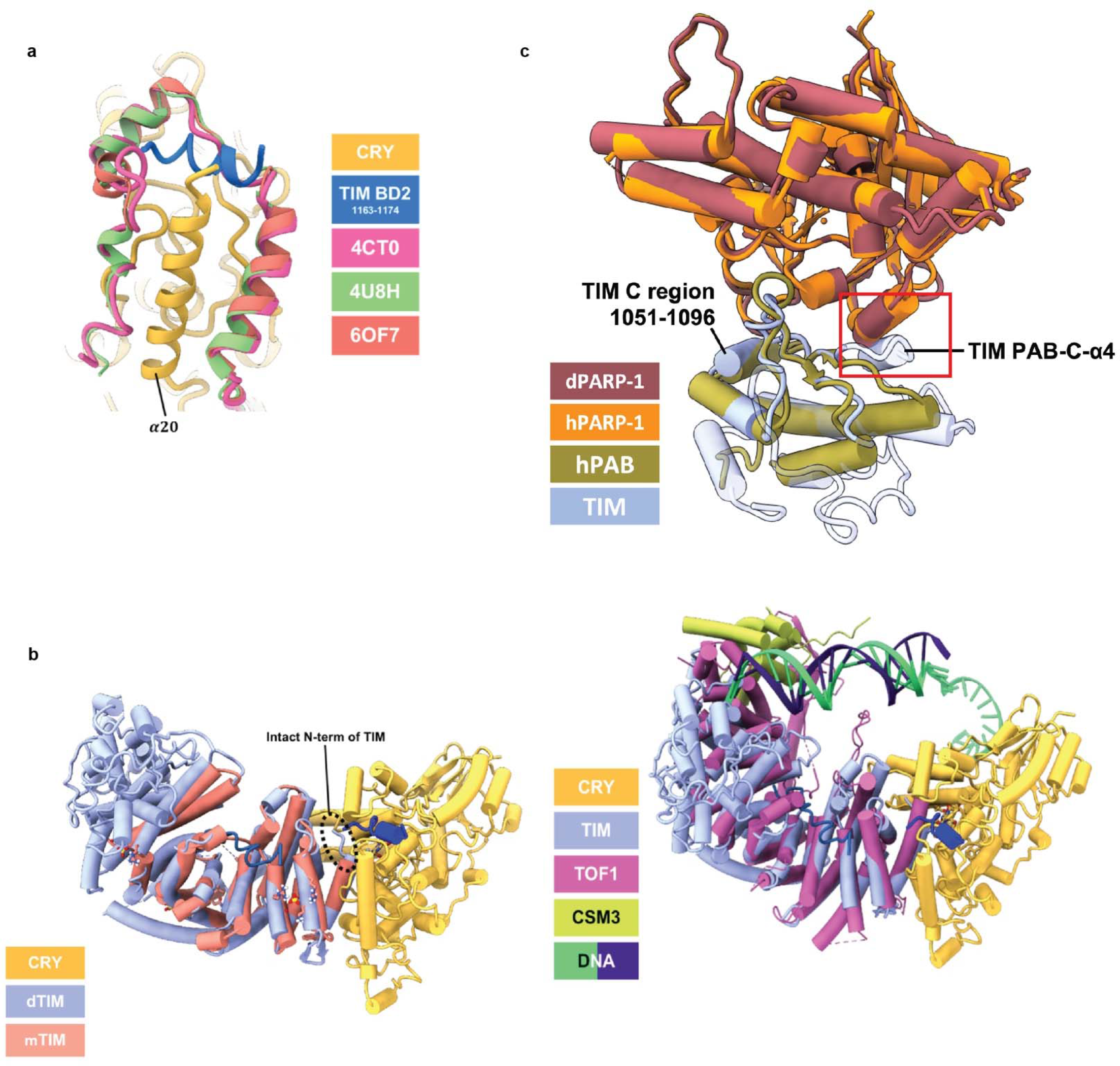
Comparisons of structural features in the CRY:TIM complex to those of CRY and TIM homologues. (a) Structural comparison of CRY:TIM with mammalian CRY bound with mammalian PER. TIM 1163-1174 (**blue**) follows the backbone trace of mPER (CRYΔ (**yellow**), TIM BD2 1163-1174 (**blue**)). 4CT0: Crystal Structure of Mouse Cryptochrome1 in Complex with Period2 (**magenta**). 4U8H: Crystal Structure of Mammalian Period- Cryptochrome Complex (**green**). 6OF7: Crystal structure of the CRY1-PER2 complex (**orange red**). (b) Structural comparison of CRY:TIM (CRY (**yellow**), TIM (**periwinkle blue**), TIM 1163- 1174 (**blue**), TIM groove peptide (**navy blue**)) with the crystal structure of the N-terminal domain of human Timeless (PDB ID: 5MQI, **salmon**). Note that mammalian TIM does not contain the N-terminal residues that bind into the CRY flavin pocket. (c) Structural superposition of TIM PAB (PARP-1 binding) domain and PARP-1 model with the structure of human TIM PAB bound to the PARP-1 catalytic domain (4XHU). Red box highlights the additional TIM α-helix (PAB-C-α4) that is incompatible with the interface formed by human PARP-1 and TIM PARP-1 binding. The TIM C region conserved across TIM proteins (residue 1051-1096) is also shown. TIM (this study, **periwinkle blue**), *Drosophila* PARP-1 (AlphaFold2.1 model: **dark red**) 4XHU (PAB:**green**, PARP-1:**orange**). (d) Superposition of TIM and Tof1 within the replisome.

**Extended Data Figure 10.**
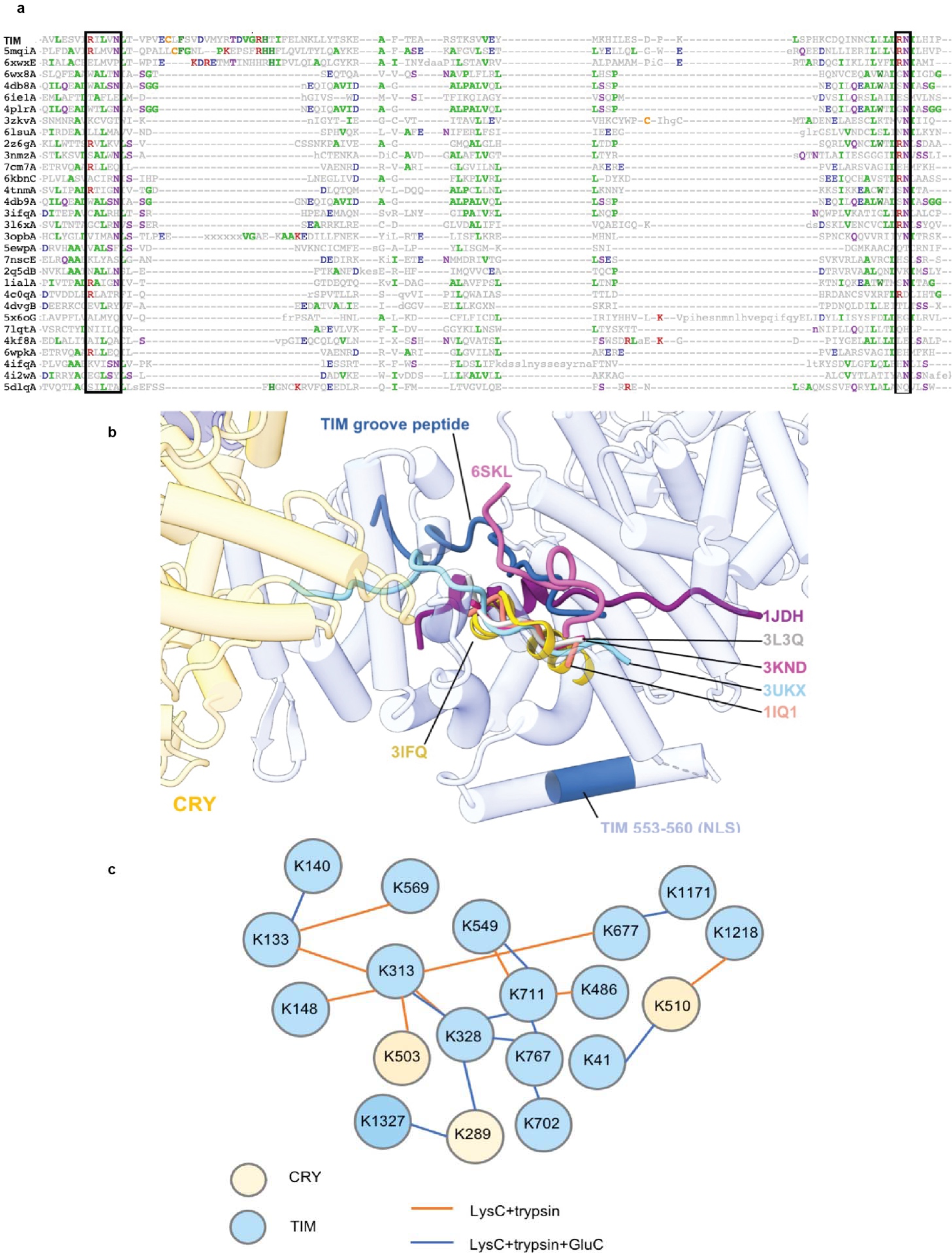
Interactions between TIM and its homologs with extended polypeptides in their respective ARM grooves in relation to crosslinking map of CRY:TIM. (a) Sequence alignment of top TIM structural homologs. Boxes outline the conserved R-XXX-N and RN residues within the ARM motifs that interact with the extended polypeptide chains of binding partners. Protein Data Bank (PDB) codes are shown on the left. (b) The extended peptide bound in the TIM groove resembles interactions found in other ARM-repeat proteins such as Importin-α1, β-catenin, and yeast TIM proteins. ARM-repeat proteins were structurally superimposed by their ARM-repeat cores to show the alignment of the extended peptides that bind in the crescent groove common to the family. For clarity only the groove peptides are displaced. Extended polypeptides found in yeast TOF1 (6SKL) and β-catenin (3IFQ) and TIM travel on the same lateral axis anti-parallel to the other importin- α, β-catenin, and TIM protein homologues. The PDB labels indicate the N-terminus of the loop. 6SKL: Cryo-EM structure of the CMG Fork Protection Complex at a replication fork - Conformation 1 (lavender), 3IFQ: Interaction of plakoglobin and β-catenin with desmosomal cadherins (**gold**), 1JDH: Crystal structure of β- catenin and htcf-4 (**purple**), 3L3Q: Mouse importin α-pepTM NLS peptide complex (gray), 3KND: TPX2:importin-αlpha complex (**magenta**), 1IQ1: Crystal structure of the importin-α(44-54)- importin-α(70-529) complex (**salmon**), 3UKX: Mouse importin-α:Bimax2 peptide complex (**sky blue**). (c) Map of DSSO lysine residue crosslinking defined by mass spectroscopy for the CRY:TIM complex. For the LysC+trypsin digested sample, although +2-charged peptides were excluded in the analysis, 59.79% sequence coverage for CRY and 49.81% for TIM were obtained. For the LysC+trypsin+GluC digested sample, 25.07% sequence coverage was obtained for CRY and 25.53% sequence coverage was obtained for TIM. Summary of the cross-links are given in the attached excel files and the raw data has been submitted to the PRIDE database.

## Supplementary Information

### Supplementary Methods

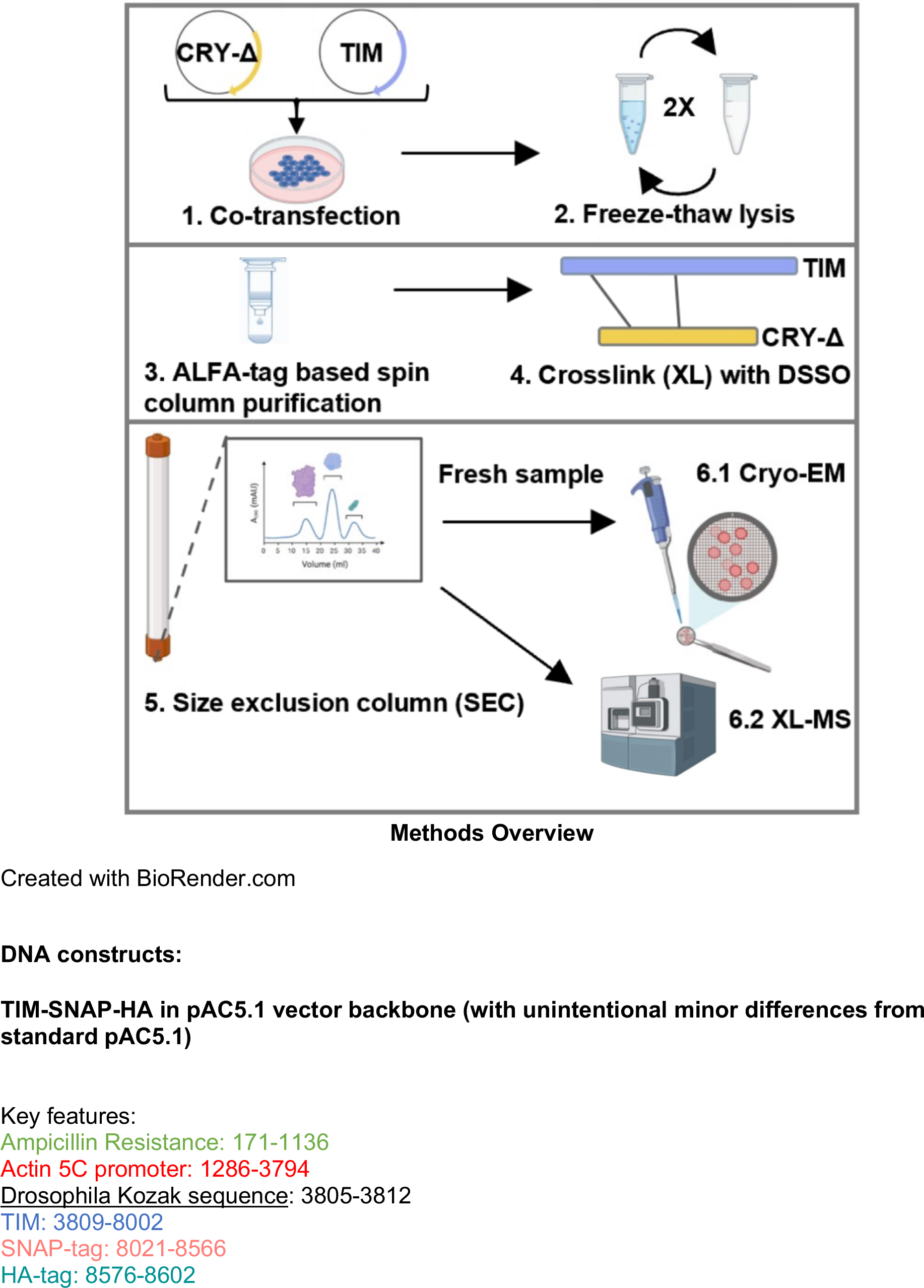
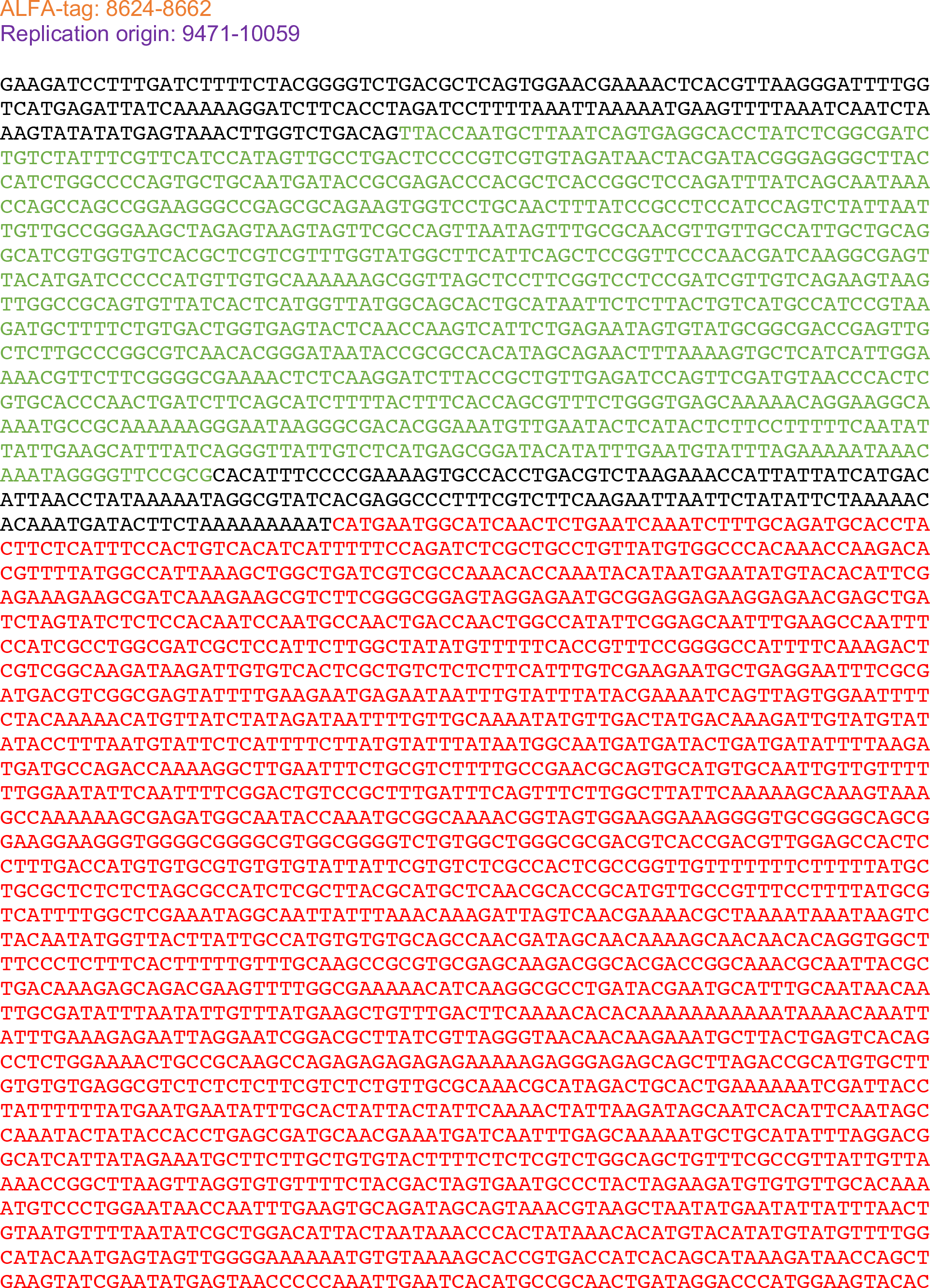
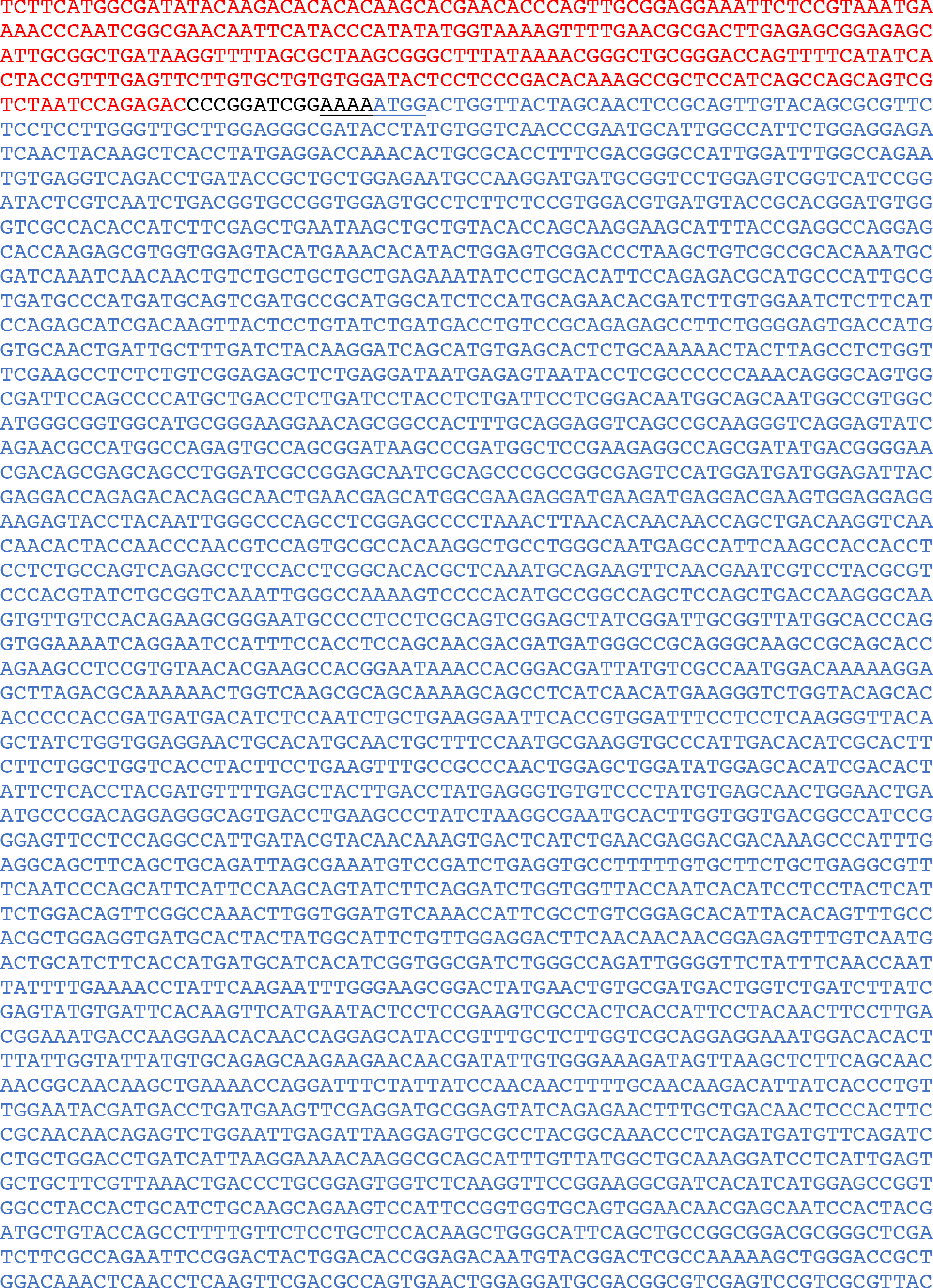
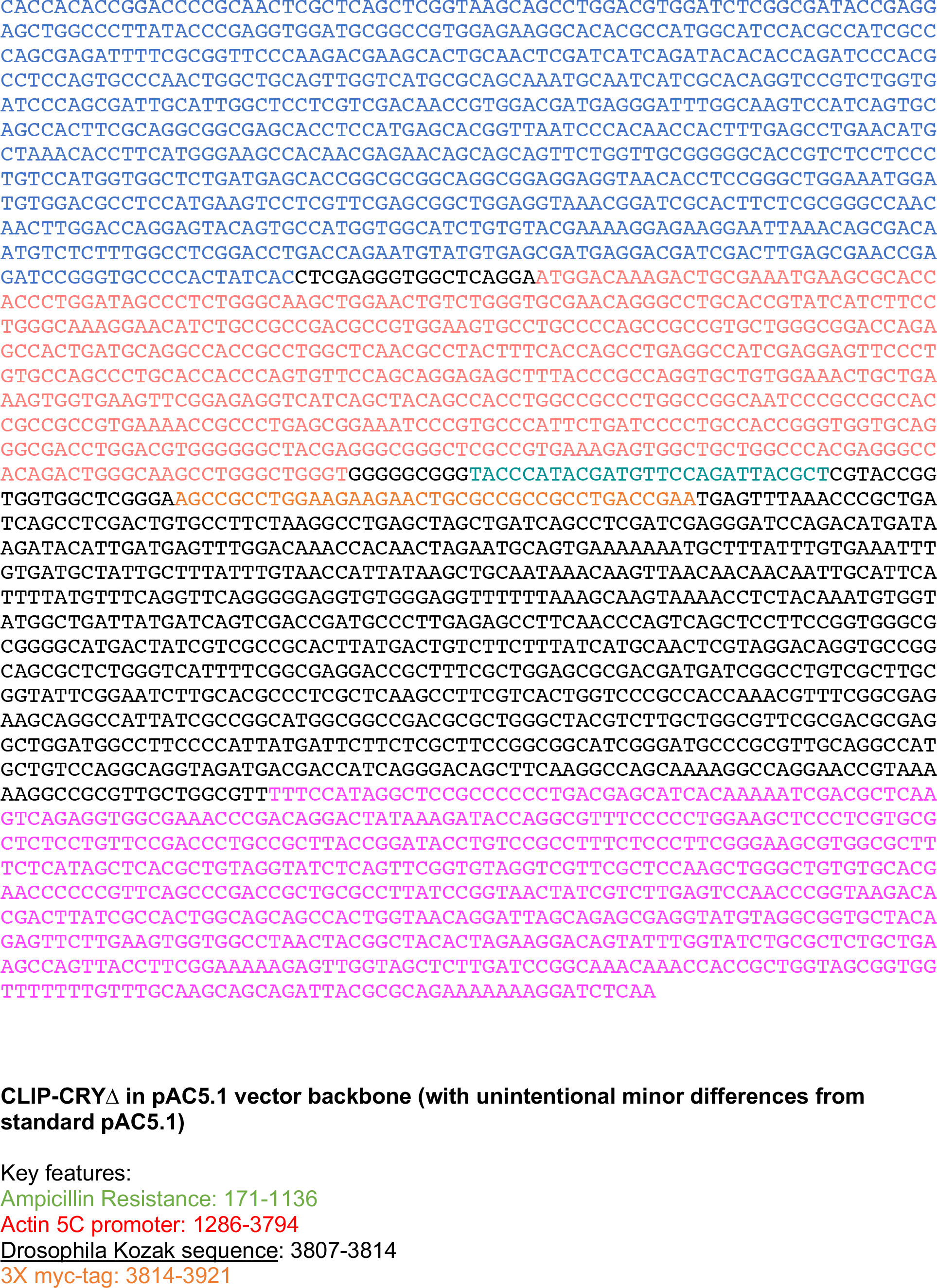
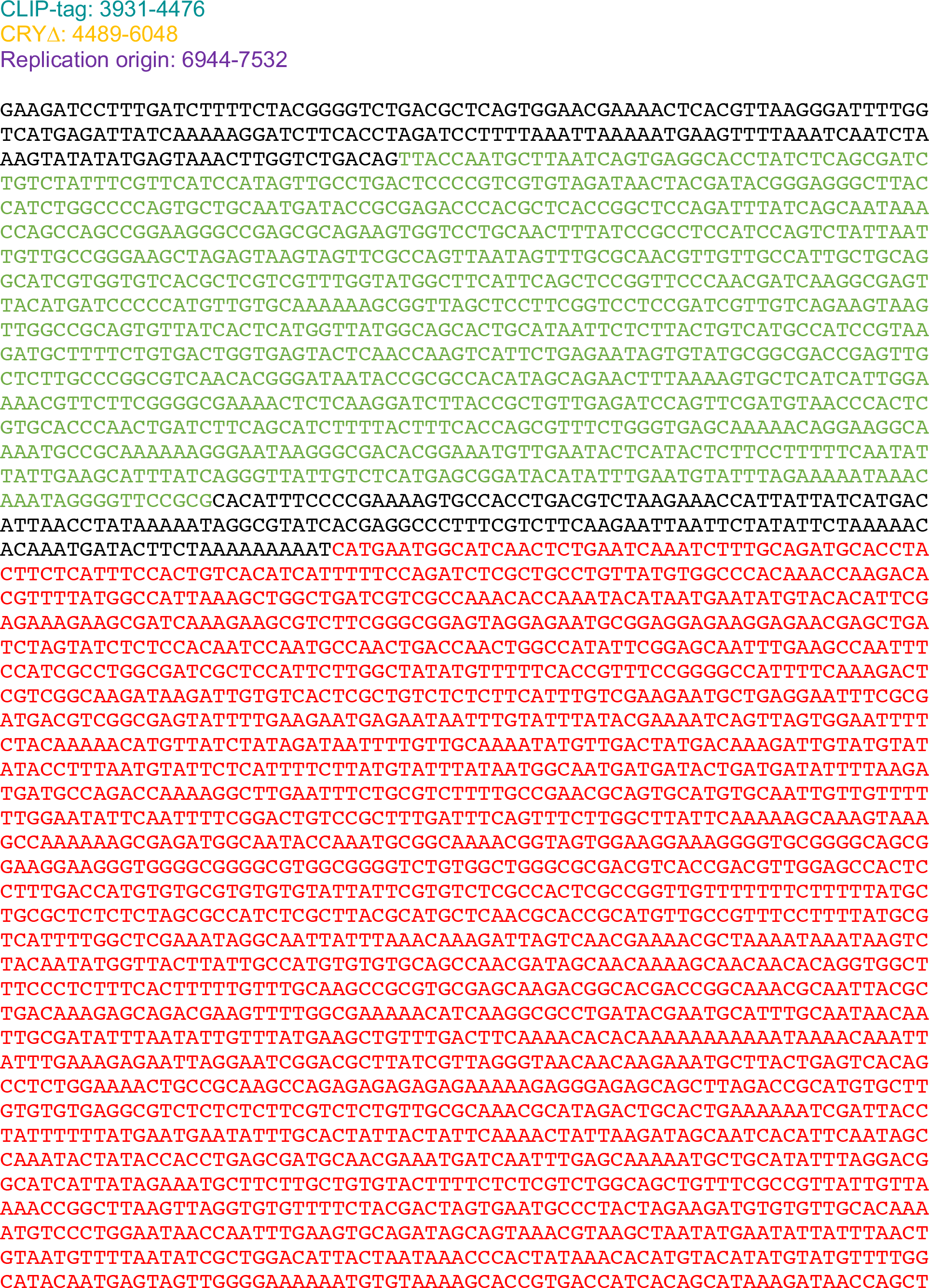
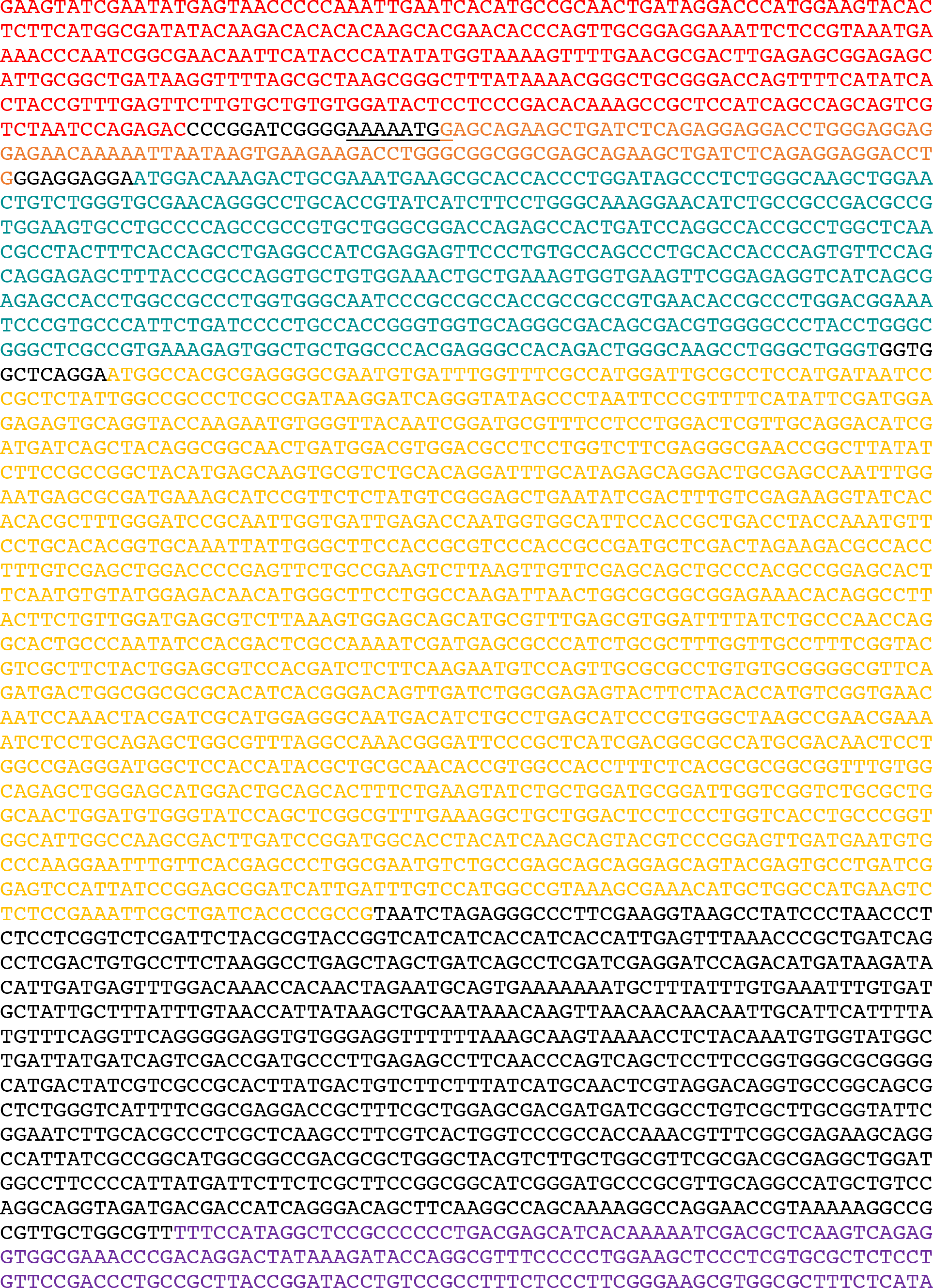
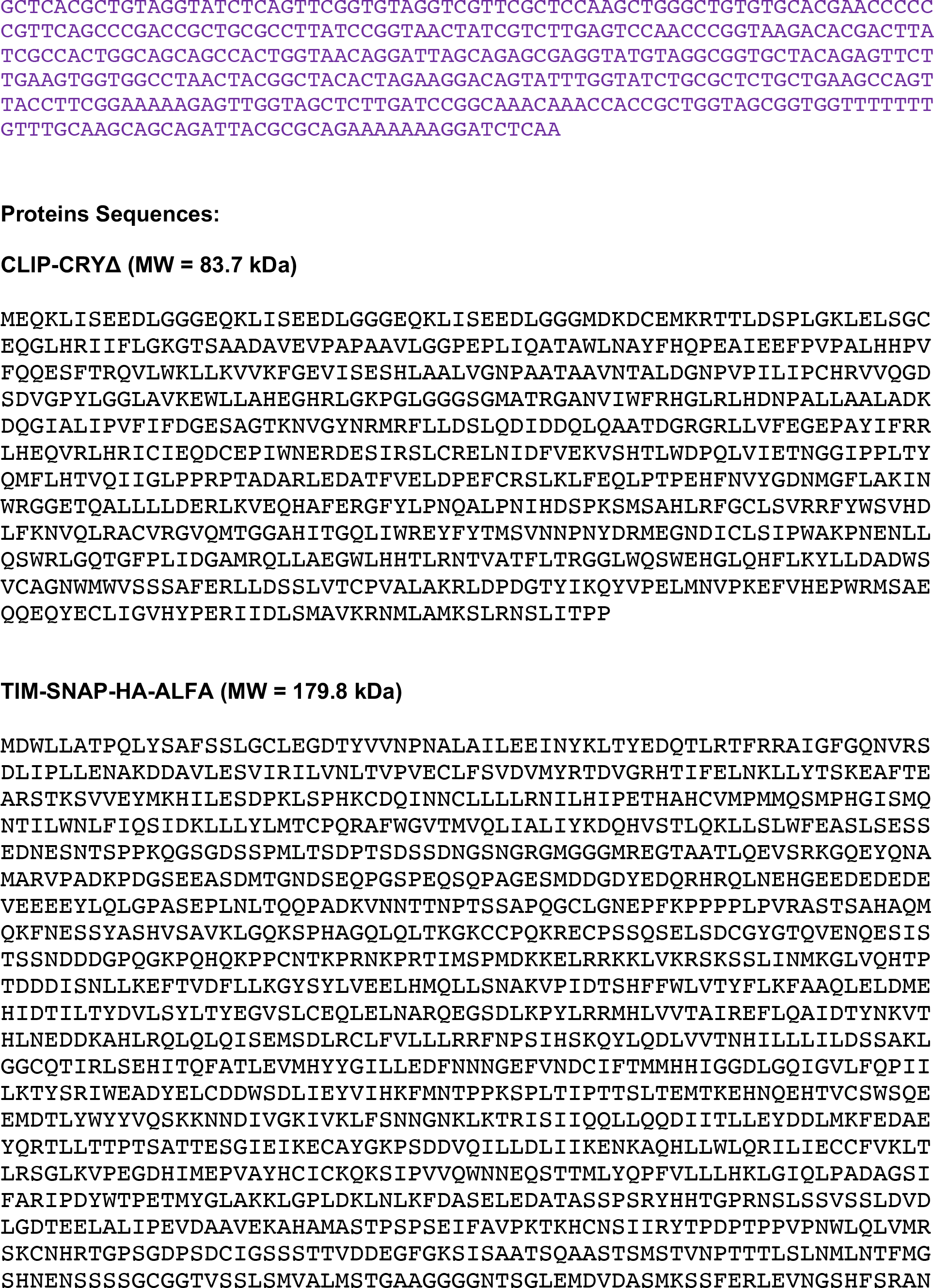
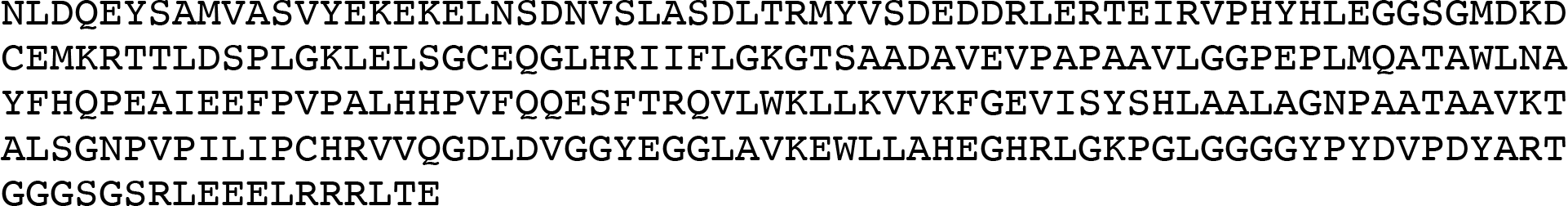

**Supplementary Table 1.**
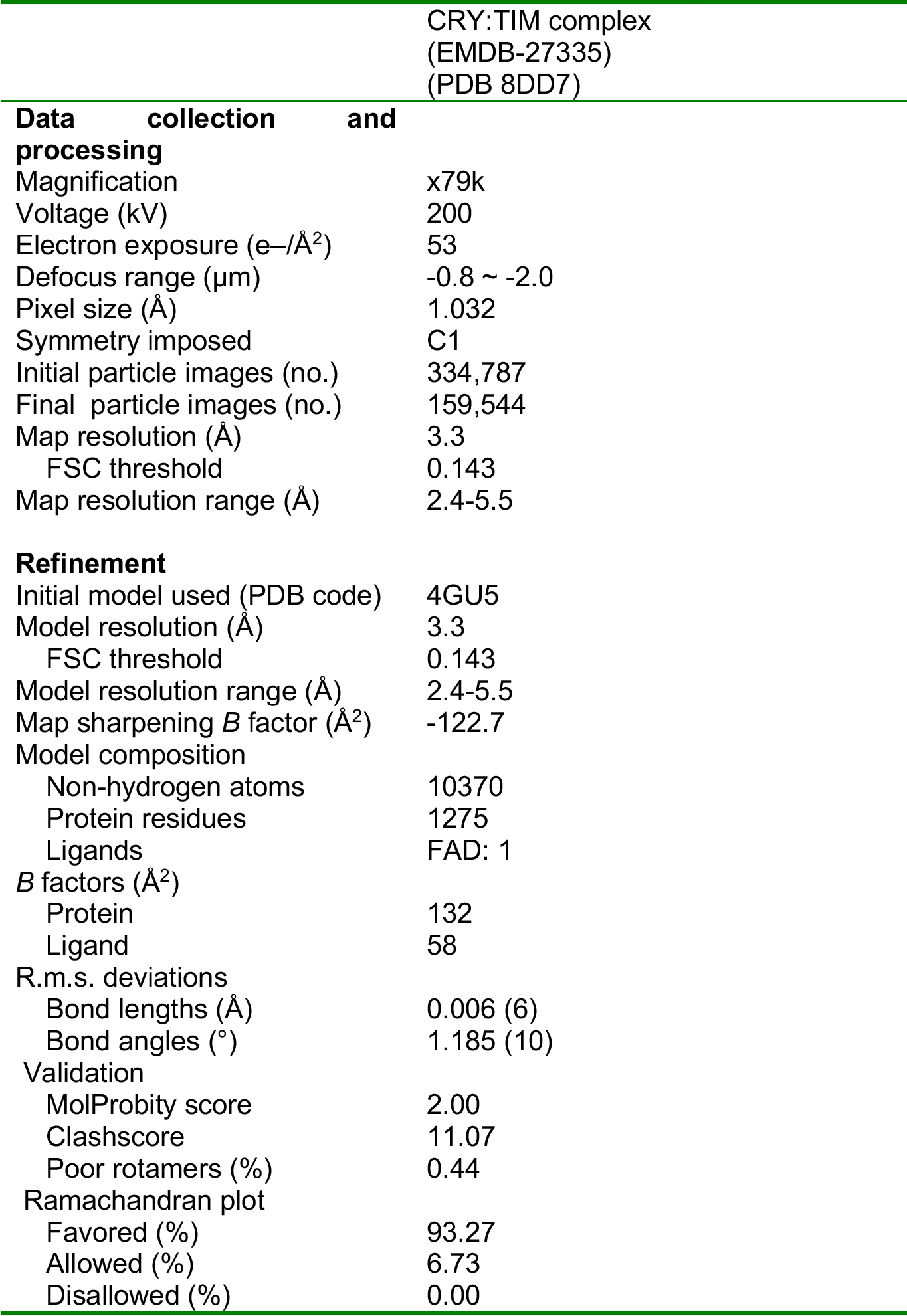
Cryo-EM data collection, refinement, and validation statistics

|  |  | CRY:TIM complex<br>(EMDB-27335)<br>(PDB 8DD7) |
| --- | --- | --- |
| **Data collection and processing** |  |  |
| Magnification |  | x79k |
| Voltage (kV) |  | 200 |
| Electron exposure (e-/Å<sup>2</sup>) |  | 53 |
| Defocus range (μm) |  | -0.8 ~ -2.0 |
| Pixel size (Å) |  | 1.032 |
| Symmetry imposed |  | C1 |
| Initial particle images (no.) |  | 334,787 |
| Final particle images (no.) |  | 159,544 |
| Map resolution (Å) |  | 3.3 |
| FSC threshold |  | 0.143 |
| Map resolution range (Å) |  | 2.4-5.5 |
| **Refinement** |  |  |
| Initial model used (PDB code) |  | 4GU5 |
| Model resolution (Å) |  | 3.3 |
| FSC threshold |  | 0.143 |
| Model resolution range (Å) |  | 2.4-5.5 |
| Map sharpening *B* factor (Å<sup>2</sup>) |  | -122.7 |
| Model composition |  |  |
| Non-hydrogen atoms |  | 10370 |
| Protein residues |  | 1275 |
| Ligands |  | FAD: 1 |
| *B* factors (Å<sup>2</sup>) |  |  |
| Protein |  | 132 |
| Ligand |  | 58 |
| R.m.s. deviations |  |  |
| Bond lengths (Å) |  | 0.006 (6) |
| Bond angles (°) |  | 1.185 (10) |
| Validation |  |  |
| MolProbity score |  | 2.00 |
| Clashscore |  | 11.07 |
| Poor rotamers (%) |  | 0.44 |
| Ramachandran plot |  |  |
| Favored (%) |  | 93.27 |
| Allowed (%) |  | 6.73 |
| Disallowed (%) |  | 0.00 |

